# Slow relaxation oscillations in multi-scale adaptive next generation neural masses

**DOI:** 10.64898/2026.07.26.740760

**Authors:** Gianluca Martelloni, David Angulo Garcia, Giacomo Innocenti, Alessandro Torcini, Simona Olmi

## Abstract

We have studied the emergence of slow relaxation oscillations in next generation neural mass models with spike frequency adaptation. Relaxation oscillations connect low firing state (Down state) to high firing state (Up state) via the slow adaptation. In the examined cases, the orbit relaxes towards the Up State via a sequence of collective damped oscillations (peaks of activity), thus revealing population bursting dynamics. The slower is the adaptation time scale the higher is the complexity (number of peaks) displayed by the relaxation oscillations. In particular, a chaos-induced spike-adding mechanism regulates the increase in the number of peaks. In analogy to what found in the Hidmarsh-Rose neuron model, two different types of chaotic behaviors have been identified: Population Spiking and Population Bursting Chaos. The increase of the adaptation strength leads to shorter (longer) Up (Down) state durations somehow mimicking the effect of charbachol in *in vitro* experiments, where spontaneous slow waves are observed. Indeed, the scenario depicted in [1], where an increase of the concentration of carbachol induces a transition from anesthesia-like to sleep-like dynamics is consistent with our results based on the variation of the adaptation strength.

**Highlights:**

- Spike Frequency Adaptation (SFA) promotes the emergence of Slow Relaxation Oscillations
- Spike-adding mechanisms, controlled by SFA, lead to Relaxation Oscillations of increasing complexity
- Two types of chaotic behaviours: Population Spiking and Population Bursting Chaos
- SFA regulates Up and Down States’ durations and their correlation

## 1. Introduction

In the context of computational neuroscience, mean-field models describing the macroscopic evolution of a homogeneous population of neurons in terms of few collective variables are referred as neural mass models [2]. These models range from heuristic firing-rate models, characterized by a single scalar variable, as the Wilson-Cowan model [3] to more refined versions based on an eigenfunction expansion of the Fokker-Planck equation ruling the evolution of the distribution of the membrane potentials of spiking neurons [4, 5]. In the last decade it has been introduced a new class of neural mass models [6, 7, 8], termed *next generation neural mass models* (NG-NMMs) [9], able to describe in an exact manner the macroscopic evolution of globally coupled heterogeneous populations of Quadratic Integrate-and-Fire (QIF) neurons[10].

The exact low-dimensional mean-field description can be obtained thanks to the fact that the QIF neuron is mathematically equivalent to the Θ-neuron, a phase oscillator model [11]. This equivalence allows for the Ott–Antonsen Ansatz [12] to be applied to spiking neuronal networks of this kind. This exact formulation has been extended to spiking networks in presence of global adaptation terms, like short-term depression and facilitation [13] and spike-frequency adaptation (SFA) [14, 15, 16]. SFA is a fatigue mechanism for which an active neuron, subject to some constant stimulus, gradually reduces its firing rate [17]. SFA promotes the emergence of collective events where neurons burst in a synchronized manner [18]. In particular, NG-NMMs can display collective events resembling spiking or bursting dynamics at the single-neuron level [14]. Population spiking (PS) correspond to a highly synchronized collective firing event, while for sufficiently long adaptation scale a population burst can be usually seen as a relaxation oscillation (RO) connecting a low firing node to a high firing focus [15].

At the neural population level, during both sleep and anesthesia, slow neural oscillations emerge from the alternation between transients of high neural activity (Up states) and transients of near silence (Down states) [19]. Adaptation has been considered crucial for modeling these *slow waves*. In [20] slow waves have been interpreted as relaxation oscillations connecting high and low firing state driven by the adaptation variable. In particular, SFA in excitatory cells generates a self-inhibitory effect that destabilizes the Up state and triggers a transition back to the Down state. A completely different modeling of slow waves is reported in [21], based on the coexistence of a high and a low firing state, where the transitions between Up and Down states are triggered by fluctuations and those from Down to Up state are due to synchronous high-amplitude events. In this context, the main role of adaptation is to induce correlations among the durations of Up and Down states.

In this work, following [20], we consider the ROs, emerging in NG-NMMs, as a simplified representation of cortical slow waves. Accordingly, we use NG-NMMs as benchmark models that retain the essential mechanisms required for the emergence of ROs, as discussed in [20, 22]. Our goal is not to reproduce all the physiological details of slow-wave activity through highly refined neural models, such as the compartmental network model reported in [23] or the biologically realistic mean-field models derived in [24]. Instead, we focus on specific statistical properties of slow waves, including the durations of Up and Down states and the correlations between consecutive states. These features have been the subject of extensive experimental investigation over the last fifteen years [25, 21, 1]; a recent overview of this line of research can be found in [26].

The remainder of the paper is organized as follows. Section 2 introduces the network model and outlines the derivation of the corresponding neural mass models. The same section also presents the linear stability analysis of both stationary and periodic solutions. Section 3 contains the main results and is divided into two parts. The first part focuses on a purely excitatory population with SFA, where we characterize the bifurcation structure of the model and investigate the dynamical mechanisms leading from PS to ROs with particular emphasis on the role played by excitability and adaptation strength. The second part considers an excitatory–inhibitory neural mass model with SFA acting only on the excitatory population. After describing the main dynamical regimes, we examine the role of inhibition in shaping relaxation oscillations of increasing complexity and discuss the qualitative correspondence between the model dynamics and experimentally observed slow waves during sleep and anesthesia. Finally, Section 4 summarizes the main findings and discusses their implications.

## 2. Methods

### 2.1. Model

We consider a fully coupled spiking neural network composed of two interacting populations, one excitatory (E) and one inhibitory (I), incorporating a biologically motivated spike-frequency adaptation mechanism that accounts for neuronal fatigue and progressively reduces neuronal excitability following spike emission. This mechanism is the so-called *spike-frequency adaptation*, which can manifest itself in different forms [17]. We have chosen to mimic this effect by following [18, 27, 28] as a slow inactivation (inhibitory) current acting only on the excitatory neurons and characterized by a time scale *τ*_*a*_. As reported in [27], pyramidal and fast spiking neurons in the rat somatosensory cortex can display both facilitation and adaptation (depression). In particular, for pyramidal neurons, adaptation is a process characterized by multiple time scales, ranging from fast adaptation with *τ*_*a*_ ≃ 50 ms to slow adaptation with *τ*_*a*_ in the range [1.4 : 15.7] sec.

We assume that each population contains an equal number *N* of QIF neurons. Therefore, the membrane potential 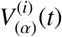 of neuron *i* belonging to the population *α* ∈ {*E, I*} evolves according to the following differential equation:

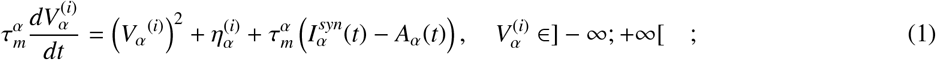

where 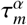 is the membrane constant of the *α*-population, 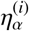 denotes the intrinsic excitability of the neuron *i* in the population *α*, drawn from a prescribed distribution *g*_*α*_(*η*). The term 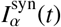 represents the total synaptic input current to the neuron *i*. Since the network is globally coupled, each neuron of population *α* is driven by the same synaptic current. *A*_*α*_(*t*) is an adaptation current, which we assume to be a global variable common to all neurons, as in [15], and acts only on excitatory ones.

Synaptic interactions are assumed to be instantaneous and are modeled as delta functions triggered by pre-synaptic spikes. The total synaptic input to any neuron in the population *α* is given by

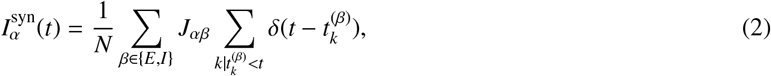

where *J*_*αβ*_ denotes the strength of the synapses connecting neurons in the population *β* to neurons in *α*-population. The variable 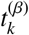 indicates the *k*-th firing time of neurons in population *β*.

As already mentioned, SFA acts only on the excitatory population, the evolution of the adaptation variable *A*(*t*) is given by

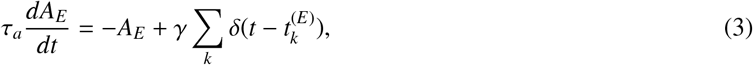

where *γ* is the adaptation strength, and the sum runs over all the spikes emitted in the excitatory population.

Neurons emit a spike when their membrane potential diverges to +∞, after which the potential is reset to −∞. This spike-and-reset mechanism is the canonical feature of the QIF model, allowing a continuous-time description of action potential generation without introducing an artificial threshold.

The first part of the paper is devoted to the study of a purely excitatory network, in this case *α* ∈ {*E}* with the remaining description left unchanged.

### 2.2. Mean-Field Description

By following [13, 15], we report the derivation of NG-NMMs with global adaptation. The neural population *α* ∈ {*E, I}* can be described in the thermodynamic limit *N* → ∞ in terms of a probability density function (PDF) *ρ*_*α*_(*V, η, t*), representing the fraction of neurons in the *α* population with membrane potential *V* and intrinsic excitability *η* at time *t*. The dynamics of *ρ*_*α*_ obey the following continuity equation:

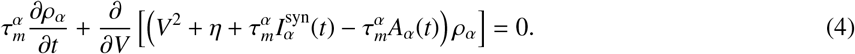

Here *A*_*α*_(*t*) is the mean-field adaptation current in population *α*, which is zero for the inhibitory population (i.e., *A*_*I*_(*t*) = 0). We assume that the intrinsic excitabilities *η* of the neurons in population *α* are distributed according to a Lorentzian distribution of the form:

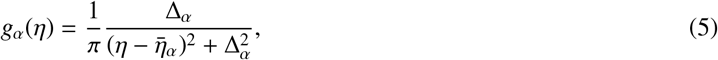

with median 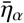 and half-width at half-maximum Δ_*α*_.

Following the approach described in [8], we look for a family of solutions of the form:

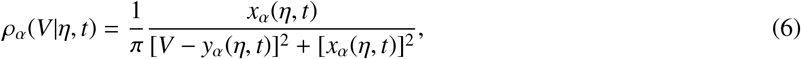

which represents a Lorentzian distribution of the membrane potentials *V*, parameterized by *x*_*α*_(*η, t*) and *y*_*α*_(*η, t*). This Ansatz provides automatically a normalization for the PDF and allows for an exact mean-field reduction of the QIF network. In particular, the macroscopic observables of interest, i.e. the population firing rate *r*_*α*_ and the mean membrane potential *v*_*α*_, are directly related to the variables appearing in (6) as follows :

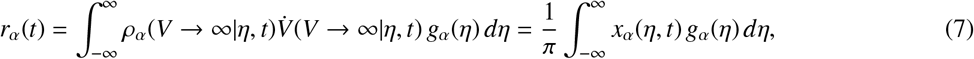

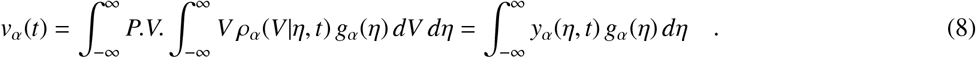

the reason for *P.V*., is that the Lorentzian distribution has a mean in the principal value sense only.

Integrals in the Eqs. (7) and (8) can be solved, once inserted the Lorentzian distribution of *g*_*α*_, by applying the Chaucy’s residue theorem. Then we can substitute the results of these integrals in the continuity equation and obtain a closed system of equations for the population firing rate *r*_*α*_ and the mean membrane potential *v*_*α*_. The result is the following set of ordinary differential equations for each population:

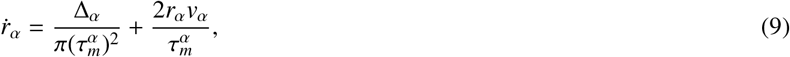

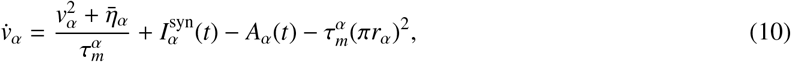

where *A*_*α*_(*t*) = 0 for *α* = *I*, and the adaptation current for the excitatory population evolves as follows:

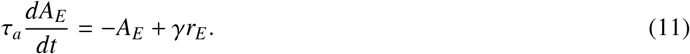

The synaptic input currents are given by:

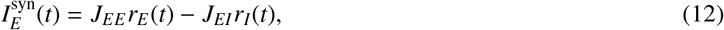

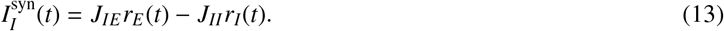

All together, this five-dimensional system of ODEs provides an exact mean-field description of the network dynamics in the limit *N* → ∞ and it captures the key dynamical features of the collective dynamics such as bifurcations among different states, level of neural synchonization, etc.

For the purely excitatory network, the mean-field description is given by Eqs. (9)-(11), with *α* = *E* and with the synaptic current given by 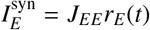.

### 2.3. Linear Stability Analysis of Fixed Points

The NG-NMM results in a five-dimensional autonomous system defined by the state variables (*r*_*E*_, *v*_*E*_, *A*_*E*_, *r*_*I*_, *v*_*I*_). The equilibrium points 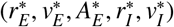, corresponding to stationary states satisfy the following set of equations :

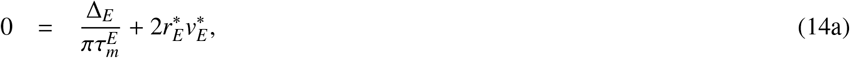

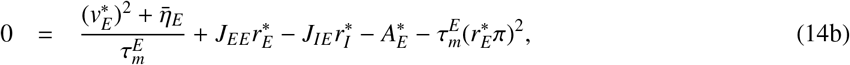

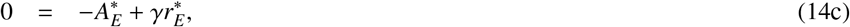

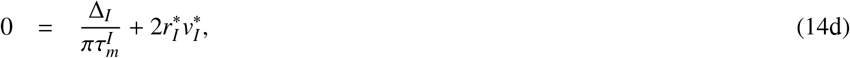

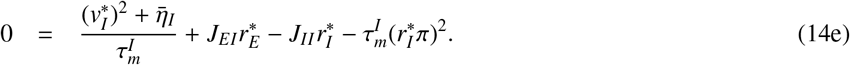

These nonlinear algebraic equations are solved numerically to find the fixed points. In general, the system may admit one or multiple equilibrium states depending on the values of the external input parameters and coupling strengths.

To analyze the local stability of each fixed point, we linearize the system around the considered equilibrium point and compute the Jacobian matrix of the system, which takes the following form:

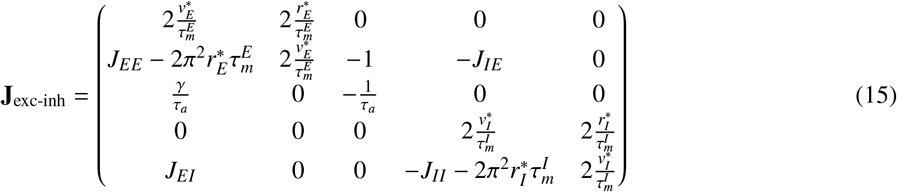

The eigenvalues of **J**_exc-inh_ determine the local stability of the fixed point. The fixed point is stable if all eigenvalues have negative real parts, whereas the presence of at least one eigenvalue with positive real part renders it unstable.

Following a similar approach for the purely excitatory network results in the following Jacobian matrix

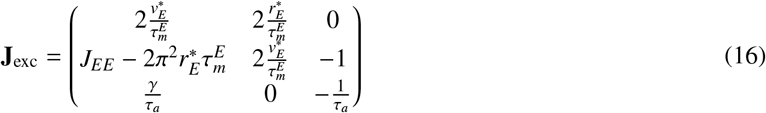

The relatively simple structure of the purely excitatory case allows us to parametrize the bifurcation structure. Dropping the subscript *E* for simplicity, the characteristic polynomial of (16) is given by

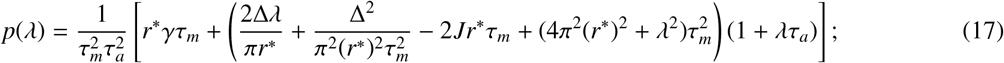

where *λ* are the -possibly-complex eigenvalues of the Jacobian, and where we made use of the equivalence *v*^∗^ = −Δ*/*(2*πr*^∗^*τ*_*m*_).

By setting *λ* = *iω* in (17) and equating it to zero, we can solve for the values where the eigenvalues of the system cross the imaginary axis. Separating real and imaginary parts of *p*(*λ* = *iω*) = 0 leads to the following expressions for the critical frequencies:

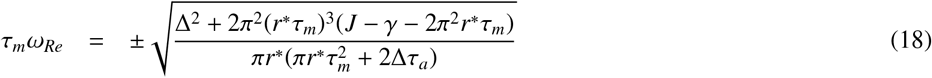

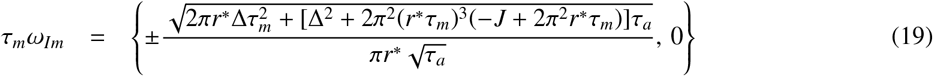

where *ω*_*Re*_ (*ω*_*Im*_) is obtained by solving Re{*p*(*iω*)} = 0 (Im{*p*(*iω*)} = 0). In each case we have two solutions differing by sign and corresponding to the two complex conjugate eigenvalues that cross the real axis, for *ω*_*Im*_ we find also the degenerate solution corresponding to zero-frequency bifurcations.

Equating the solutions in (18) and the zero solution in (19) and solving for one parameter gives the parametrization of the zero-frequency bifurcations in terms of *r*^∗^ for that parameter. Similarly, combining the non-zero solution of Eq. (19) with either branch of Eq. (18) provides a parametrization of the Hopf bifurcations. For illustrative purposes, we choose *γ* as the target parameter and we focus on the Hopf solutions. This leads to the expression

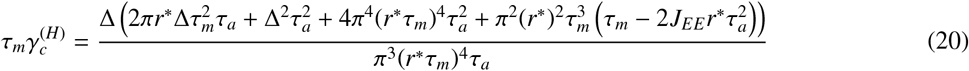

Finally, plugging the expression for 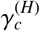 into the equilibrium expression for 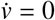 we can solve for another target parameter, say for instance 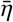 leading to the critical values of 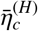 where the Hopf bifurcation occurs.

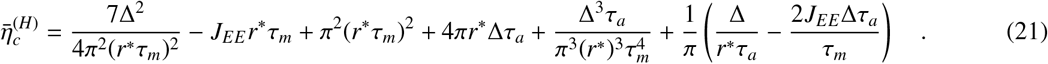

A similar procedure with the zero-frequency solutions -typically fold bifurcations-leads to:

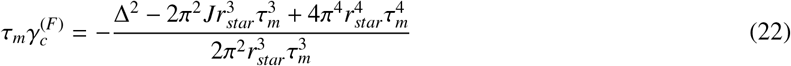

and

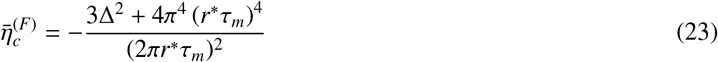

### 2.4. Linear Stability of Periodic Orbits

To assess the stability of the periodic solutions of the mean-field system, we compute the Floquet multipliers associated with each periodic orbit. Let *y*^(*p*)^(*t*) be a periodic solution of period *T*, satisfying *y*^(*p*)^(*t* + *T*) = *y*^(*p*)^(*t*) for all *t*. To analyze its linear stability, we introduce a small perturbation *ξ*(*t*) such that *y*(*t*) = *y*^(*p*)^(*t*) + *ξ*(*t*). Linearization around the periodic orbit yields the variational equation

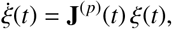

where **J**^(*p*)^(*t*) = *Df* (*y*^(*p*)^(*t*)) is the Jacobian matrix of the vector field evaluated along the orbit. In the case of the excitatory only network 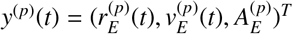 and the time dependent Jacobian is

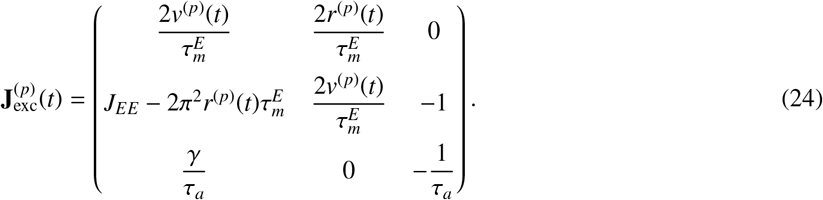

Meanwhile, for the excitatory-inhibitory network 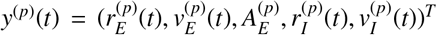 and its associated linearized evolution reads as:

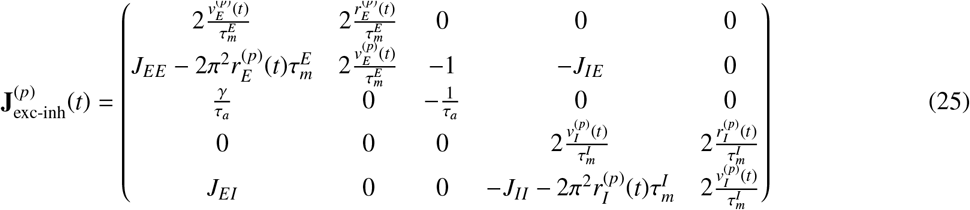

The evolution of perturbations *ξ*(*t*) over one period is governed by the *monodromy matrix* Φ(*T*), defined as the solution of the matrix differential equation

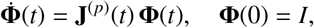

where *I* is the 3 × 3 (5 × 5) identity matrix for the purely excitatory (excitatory-inhibitory) network. The columns of Φ(*T*) represent the images of the canonical basis vectors after one full period of the linearized flow.

The eigenvalues of Φ(*T*), denoted by *µ*_*i*_, are the *Floquet multipliers*. These quantify how infinitesimal perturbations evolve after one cycle of the periodic orbit. If all multipliers satisfy |*µ*_*i*_| *<* 1, except for the trivial one *µ*_1_ = 1 (associated with time invariance), the periodic solution is asymptotically stable. Conversely, if one or more multipliers satisfy |*µ*_*i*_| *>* 1, the orbit is unstable.

The crossing of the value one of the modulus of the Floquet multipliers determine the onset of different types of bifurcations of periodic orbits, namely:

1. When a real multiplier crosses +1, the system undergoes a *saddle-node (fold)* of limit cycles.
2. When a real multiplier crosses −1, a *period-doubling (flip)* bifurcation occurs.
3. When a complex-conjugate pair of multipliers crosses the unit circle (|*µ*_*i*_| = 1), a *Neimark–Sacker (torus)* bifurcation arises.

### 2.5. Lyapunov Exponents

To analyze the nature of the macroscopic solutions of Eqs. (9-11), we estimate the corresponding Lyapunov spectrum (LS) [29]. This is achieved by considering the time evolution of the tangent vector, which for the excitatory-inhibitory set-up turns out to be five dimensional, i.e., *δ* = (*δr*_*E*_, *δv*_*E*_, *δA*_*E*_, *δr*_*I*_, *δv*_*I*_)^*T*^ . The dynamics of the tangent vector is ruled by the linearization of the Equations 9-11, namely

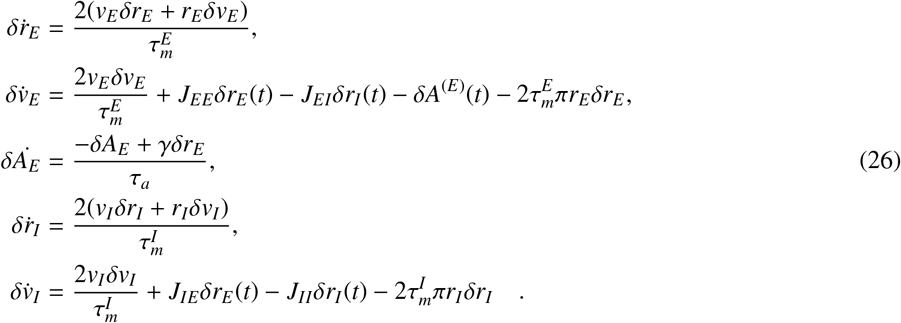

The LS consists of 5 Lyapunov exponents (LEs) {*λ*_*i*_*}* for the excitatory-inhibitory set-up, which quantify the average growth rates of infinitesimal perturbations along orthogonal manifolds. For the purely excitatory network, the LS reduces to the evolution of the first 3 equations of the system (26) and the spectrum consists of 3 LEs {*λ*_*i*_*}*.

In detail, LEs are estimated as follows:

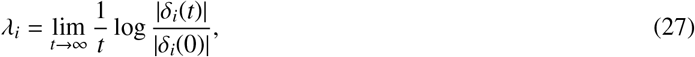

where the technique described in [30] to maintain the tangent vectors *δ*_*i*_ orthonormal during the evolution is employed. The autonomous system will be chaotic for *λ*_1_ *>* 0, while a periodic (quasi-periodic) dynamics will be characterized by *λ*_1_ = 0 (*λ*_1_ = *λ*_2_ = 0) and a fixed point by *λ*_1_ *<* 0. In a non-autonomous system in the presence of an external forcing, one Lyapunov exponent will be necessarily zero, therefore a periodic behavior corresponds to *λ*_1_ *<* 0 and a quasi-periodic dynamics to *λ*_1_ = 0 [29].

### 2.6. Kaplan–Yorke Dimension

In order to further characterize the chaotic macroscopic solutions, we estimated the Kaplan–Yorke dimension *D*_*KY*_ from the Lyapunov spectrum. This quantity provides an estimate of the effective fractal dimension of the attractor by measuring how many expanding and weakly contracting directions are dynamically involved in the long-term evolution of the system [31, 32].

Let the Lyapunov exponents be ordered in decreasing order, *λ*_1_ ≥ *λ*_2_ ≥ · · · ≥ *λ*_*N*_, where *N* = 5 for the excitatory-inhibitory set-up and *N* = 3 for the purely excitatory network. The Kaplan–Yorke dimension is defined as

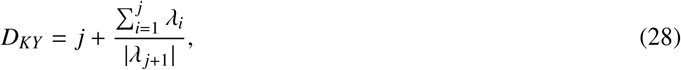

where *j* is the largest integer such that

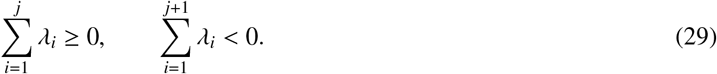

Thus, *D*_*KY*_ corresponds to an interpolation between the number of directions whose cumulative expansion is non-negative and the next contracting direction required to compensate this expansion.

For fixed-point or periodic solutions, the Kaplan–Yorke dimension remains integer or close to an integer value compatible with the corresponding low-dimensional invariant set. In contrast, for chaotic solutions with *λ*_1_ *>* 0, *D*_*KY*_ typically takes a non-integer value, reflecting the fractal geometry of the strange attractor. In the present work, *D*_*KY*_ was computed from the asymptotic Lyapunov spectrum obtained with the same orthonormalization procedure described above. Therefore, for each set of parameters, the reported Kaplan–Yorke dimension was evaluated only after discarding the initial transient and after verifying the convergence of the Lyapunov exponents.

### 2.7. Numerical Simulations and Bifurcation Analysis

Microscopic network simulations of purely excitatory neurons, described by Eqs. (1)–(3), were performed using *N*_*E*_ = 10^5^ QIF neurons over a simulation time of *t* = 2s. Time integration was carried out with a fixed-step Runge–Kutta scheme using a step size *h* = 2 × 10^−2^s. Spikes were detected using an effective threshold *V*_th_ = 100, after which the membrane potential was reset to *V*_reset_ = −*V*_th_.

Simulations of the mean-field equations were performed using an Adams/BDF method with automatic stiffness detection, as implemented in the `scipy.integrate` module. This approach ensures accurate resolution of fast transients associated with the dynamics of bursting.

Continuation of fixed points and their stability was carried out using the `MATCONT 7p3` package in `MATLAB R2022b`.

## 3. Results

### 3.1. Purely Excitatory Network with SFA

We begin by analyzing the purely excitatory network with SFA and in order to simplify the notation, the subscript or superscript E in the variables and parameters will be dropped. Let us start by comparing the network dynamics and the neural-mass model results, these are reported in Figure 1 for two distinct oscillatory dynamics, previously introduced : Population Spiking and Population Bursting [15]. Remarkably, the mean-field equations reproduce almost exactly the microscopic dynamics, validating the performed reduction even in the strongly nonlinear bursting regime. The small discrepancies observed at long simulation times are attributed to finite-size fluctuations in the microscopic network.

**Figure 1:**
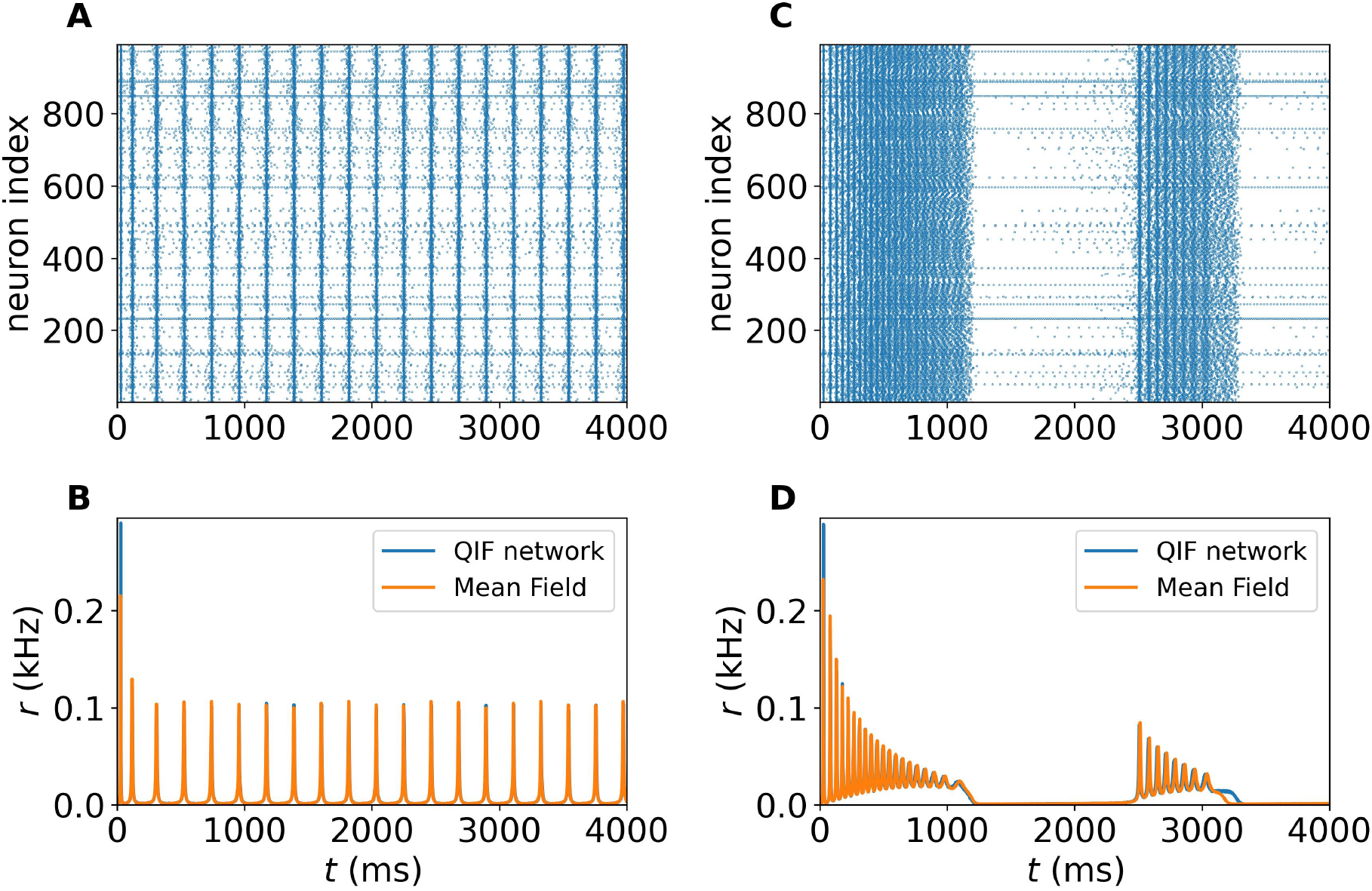
Network Simulations vs Neural Mass Evolution. (A) Raster plot in the PS regime. (B) Population firing rate *r*(*t*) obtained via microscopic simulations of the network and of the corresponding neural mass model. (C) and (D) same as in (A) and (B) for the slow ROs joined to PBs. The set of parameters used for this figure were: Δ = 0.1, *τ*_*m*_ = 20 ms, *J* = 7, *γ* = 10, 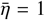. For the spiking behavior *τ*_*a*_ = 200 ms while for the slow RO *τ*_*a*_ = 200 ms. The microscopic network was simulated for *N* = 100, 000 neurons by employing an Euler integration scheme using a time step *h* = 1 × 10^−3^.

A key control parameter for these dynamics is the adaptation time constant *τ*_*a*_. For relatively short *τ*_*a*_ values, the system exhibits regular oscillations of the firing rate (referred as PS), characterized by a single time scale (Fig. 1(A-B)). In contrast, for sufficiently large *τ*_*a*_, the excitatory population undergoes bursting activity (Fig. 1(C-D)). In this regime, episodes of rapid spiking (also associated to *Up states* in analogy with Slow Wave Sleep dynamics observed in the brain) alternate with low firing periods (*Down states*), reflecting the slow recovery of adaptation. Thus, we find that the adaptation time scale *τ*_*a*_ controls the overall period of these slow relaxation oscillations (referred as ROs), while faster population bursts emerge on top of the Up state.

These results highlight two points. First, the mean-field formulation provides a reliable description of collective excitatory dynamics, in both regimes. Second, the adaptation time scale plays a critical role in shaping emergent network activity: while short *τ*_*a*_ supports regular oscillations (PSs), long *τ*_*a*_ generates slow ROs together with bursting activity by inducing a slow–fast timescale separation. These findings motivate the subsequent bifurcation analysis to clarify the mechanisms underlying the emergence of ROs in our minimal neural mass model.

We begin by analyzing the neural mass model in the absence of SFA (*γ* = 0). In this regime, the system exhibits only fixed-point solutions, as shown in Fig. 2(A). These solutions are characterized by low population firing rates for sufficiently negative values of 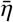 and high firing rates for sufficiently positive values. As previously shown in [8], the low activity state (LAS) is a node, while the high one (HAS) is a focus. The transition from HAS (LAS) to LAS (HAS) occurs via a saddle-node bifurcation denoted as LP_1_ (LP_2_) in panel (A) and as a consequence the two stable states are connected by an unstable saddle. As shown in panel (A), the three fixed points solution coexists in an interval of slightly negative values of 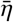.

**Figure 2:**
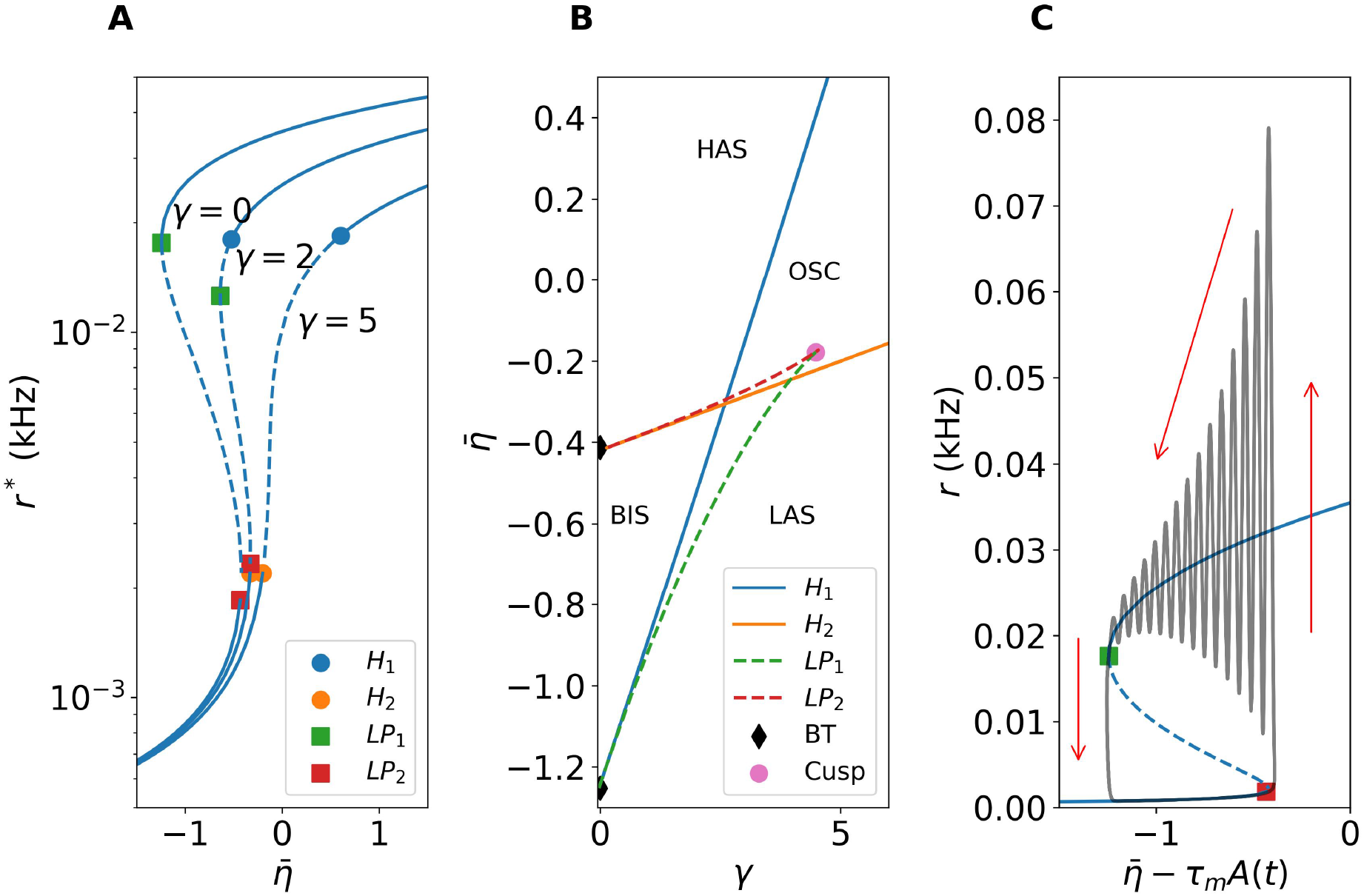
Bifurcations of High and Low Firing States. (A) Stable and unstable fixed points reported in the 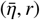-plane for three different values of the adaptation strength *γ* indicated next to each curve. Solid (dashed) lines refer to stable (unstable) fixed points. At *γ* = 0 two saddle nodes associated with high (LP_1_) and low firing rate (LP_2_) are the only two bifurcations giving rise to coexistence of stable and unstable equilibrium branches. Hopf bifurcation points associated with high (low) firing rate are indicated as H_1_ (H_2_). (B) Bidimensional bifurcation diagram in the 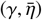-plane. Solid lines correspond to Hopf bifurcation lines, while dashed ones to Limit Point bifurcation lines with color code associated with the High/Low activity of the bifurcating stationary points. BN denotes a Bodganov-Takens bifurcation point and Cusp the cuspidal point 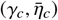. Regions with high (low) activity states are denoted with HAS (LAS) whereas the region with collective oscillations is denoted with OSC. The bistable lower triangular region where both HAS and LAS coexist is indicated with BIS. (C) Relaxation oscillation for *γ* = 5 and 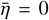 in the 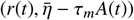 space -gray-joined to the stationary solutions of the fast sub-system - blue. We fixed *τ*_*A*_ = 2000ms, all the other parameters are as in Fig. 1.

As soon as *γ* becomes positive, the saddle-node bifurcation points LP_1_ and LP_2_ unfold into curves of Hopf and saddle-node bifurcations in the 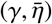 plane through two Bogdanov–Takens codimension-two points (Figs. 2(A-B)). For a fixed value of *γ >* 0, decreasing (increasing) 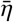 causes the HAS (LAS) to lose stability first through a Hopf bifurcation H_1_ (H_2_) and subsequently through a saddle-node bifurcation LP_1_ (LP_2_). The two saddle-node curves approach each other and eventually annihilate at the cusp point 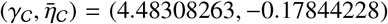. As a consequence, for *γ > γ*_C_ only the Hopf bifurcation curves remain. These Hopf bifurcation curves bound a region of collective oscillations at intermediate values of 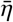 (within the upper triangular region in Fig. 2(B)), while stable HASs and LASs persist for sufficiently large positive and negative values of 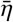, and they coexist in the lower traingular region in Fig. 2(B). The phase diagram reported in Fig. 2(B) is remarkably similar to the one reported for Leaky Integrate-and-Fire (LIF) neurons with adaptation (see [20]).

As shown in Fig. 2(C), for *τ*_*a*_ = 2000 ms a typical slow RO consists of a gradual relaxation from the HAS to the LAS, accompanied by damped collective oscillations. In this figure, as customary for the slow-fast analysis, we have reported the stable (solid blue line) and unstable equilibria (dashed blue line) of the fast sub-system (FS) (corresponding to *γ* = 0) in the plane 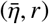 together with the oscillating solution (gray line) *r*(*t*) versus 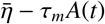 for *γ* = 5 [15]. The trajectory initially evolves along the low-firing-rate branch of the FS up to the saddle-node bifurcation point LP_2_. Beyond this point, it is attracted toward the high-firing-rate branch of the FS and approaches it through damped oscillations, reflecting the stable-focus character of the corresponding FS equilibrium. The peak of activity steadily decreases as it approaches the bifurcation point LP_1_ of the FS and, once it reaches it, the orbit jumps rapidly into the low firing manifold. In contrast to the Up state, the approach toward the low-firing-rate state is monotonic, since the corresponding equilibrium of the FS is a stable node. The durations of both the Up and Down states are primarily controlled by the adaptation timescale *τ*_*a*_. For the parameters considered here, the orbit has a period of approximately 2*τ*_*a*_ ≃ 4 s, with the Down and Up states lasting about 1.4*τ*_*a*_ and 0.6*τ*_*a*_, respectively.

Initially the Up state is characterized by damped oscillations in the population activity, with a mean frequency around 16 Hz, lying within the *β* band (13–30 Hz). As the trajectory approaches the saddle-node point LP_1_, the oscillation frequency gradually decreases from approximately 16 Hz to 8 Hz. This behavior is consistent with the properties of the corresponding stable focus of the FS. Indeed, over the interval 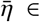 [−1.21, −0.41], -where the bursting phase exist-, the frequency associated with the damped oscillations of the stable focus decreases from about 20 Hz to 8 Hz. This confirms that the bursting oscillations are related to the damped oscillations towards the focus in the FS. Furthermore, the decrease in the oscillation amplitude and frequency is joined to a reduction of the population firing rate of the HAS as evident from Fig. 2 (C).

As a following step we considered the emergence of oscillatory solutions in the 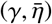-plane for different values of the adaptation time scale. To do so, we have identified the bifurcations of the equilibrium branches of the mean-field equations for three different adaptation time scales: namely, *τ*_*a*_ = {200, 500, 2000} ms. The results, shown in the top panel of Fig. 3 for 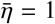, confirm that the equilibrium branch is independent of *τ*_*a*_, consistent with Eqs. (14). However, the location of the bifurcation points along this branch depends strongly on *τ*_*a*_. In all cases, the only relevant bifurcations in the parameter region of interest are two Hopf bifurcations (see colored symbols), which delimit the onset and offset of oscillatory activity.

**Figure 3:**
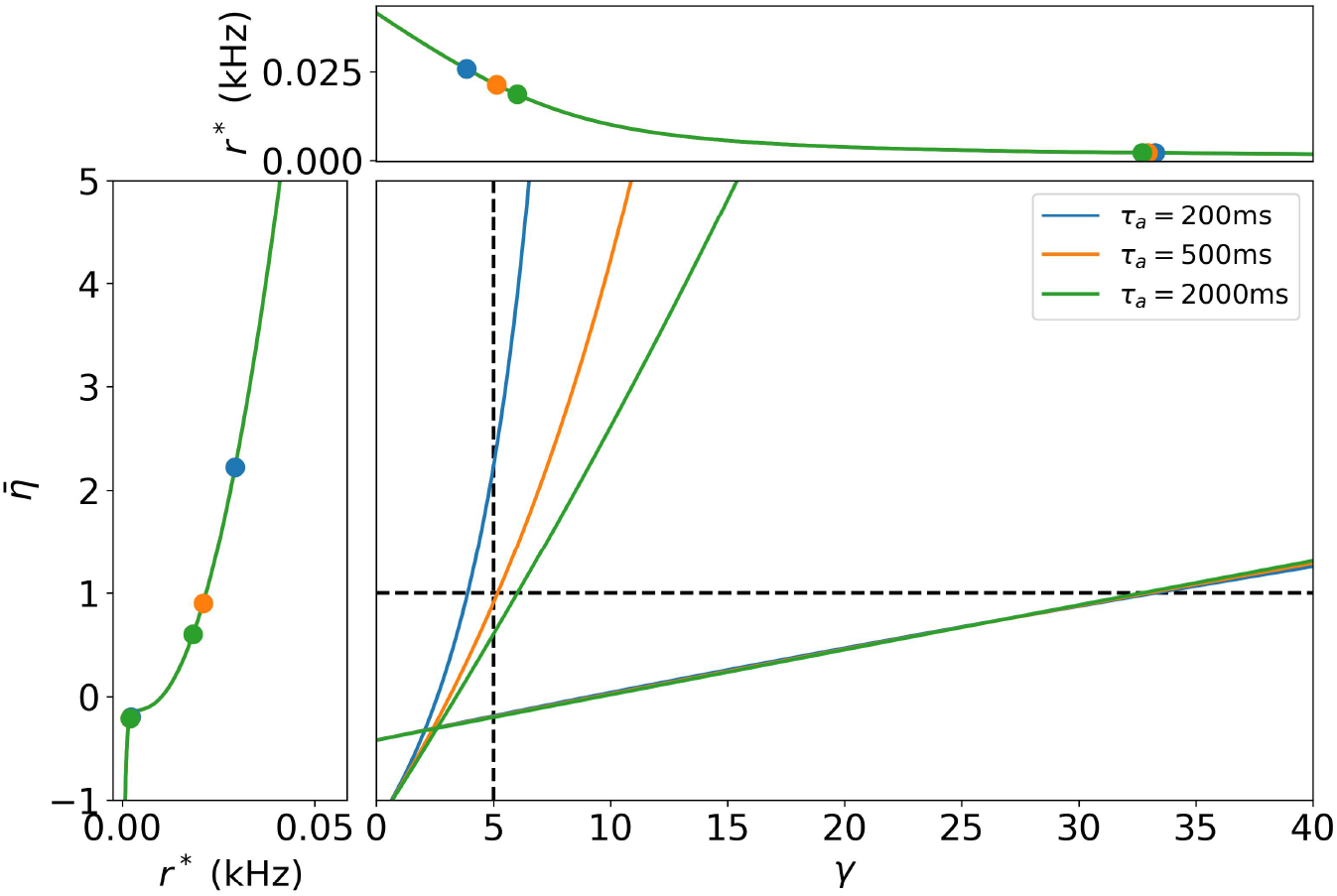
Hopf bifurcation lines. Top panel: Continuation of the equilibrium (*r* coordinate) as a function of *γ* using a fixed value of 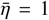 for three values of *τ*_*a*_ indicated in the legend. Left panel: Same as in top panel by fixing *γ* = 5 and varying 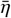. Main panel: Hopf Curves in the 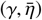 parameter plane at different *τ*_*a*_. The fixed values used for the top and left panels are shown as horizontal and vertical lines, respectively. Other parameters as in Fig. 1

A complementary analysis was performed by varying the mean excitability 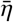 while keeping *γ* = 5 fixed (Fig. 3, left panel). For all considered values of *τ*_*a*_, the system undergoes two Hopf bifurcations. The onset of oscillatory dynamics occurs at nearly the same, slightly negative value of 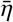, largely independent of *τ*_*a*_. In contrast, the extent of the oscillatory region depends markedly on the adaptation timescale: as *τ*_*a*_ increases, the range of 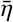 over which oscillations exist progressively shrinks.

Finally, leveraging the analytical conditions for the Hopf bifurcations derived in Eqs. (20) and (21), we constructed a codimension-2 bifurcation diagram in the 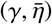-plane (Fig. 3, main panel). The diagram reveals that oscillatory activity emerges exclusively within the region enclosed by the two Hopf curves: namely, in the upper triangular region. While in the lower triangular region we have the coexistence of LAS and HAS steady solutions, -analogously to what reported in Fig. 2 (B). Furthermore, while the lower Hopf line is almost insensitive to the value of *τ*_*a*_, the upper one strongly depends on *τ*_*a*_. In summary, Fig. 3 establishes that *τ*_*a*_ does not alter the fixed-point branch itself, but rather reshapes the bifurcation landscape, modulating the extent of the oscillatory region : longer adaptation times strongly reduce the domain of the existence of oscillations.

Although the bifurcation diagrams in Fig. 3 share a similar overall structure, the dynamics inside the oscillatory region is profoundly influenced by the value of *τ*_*a*_. To characterize these orbits, we estimated the local maxima *r*_*M*_ of the population firing rate *r* versus *τ*_*a*_ for fixed values of 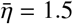 and *γ* = 8 by performing quasi-adiabatic simulations, where *τ*_*a*_ is increased/decreased in steps Δ*τ*_*a*_. ^2^ The corresponding values of *r*_*M*_ are shown in Fig. 4 (A).

**Figure 4:**
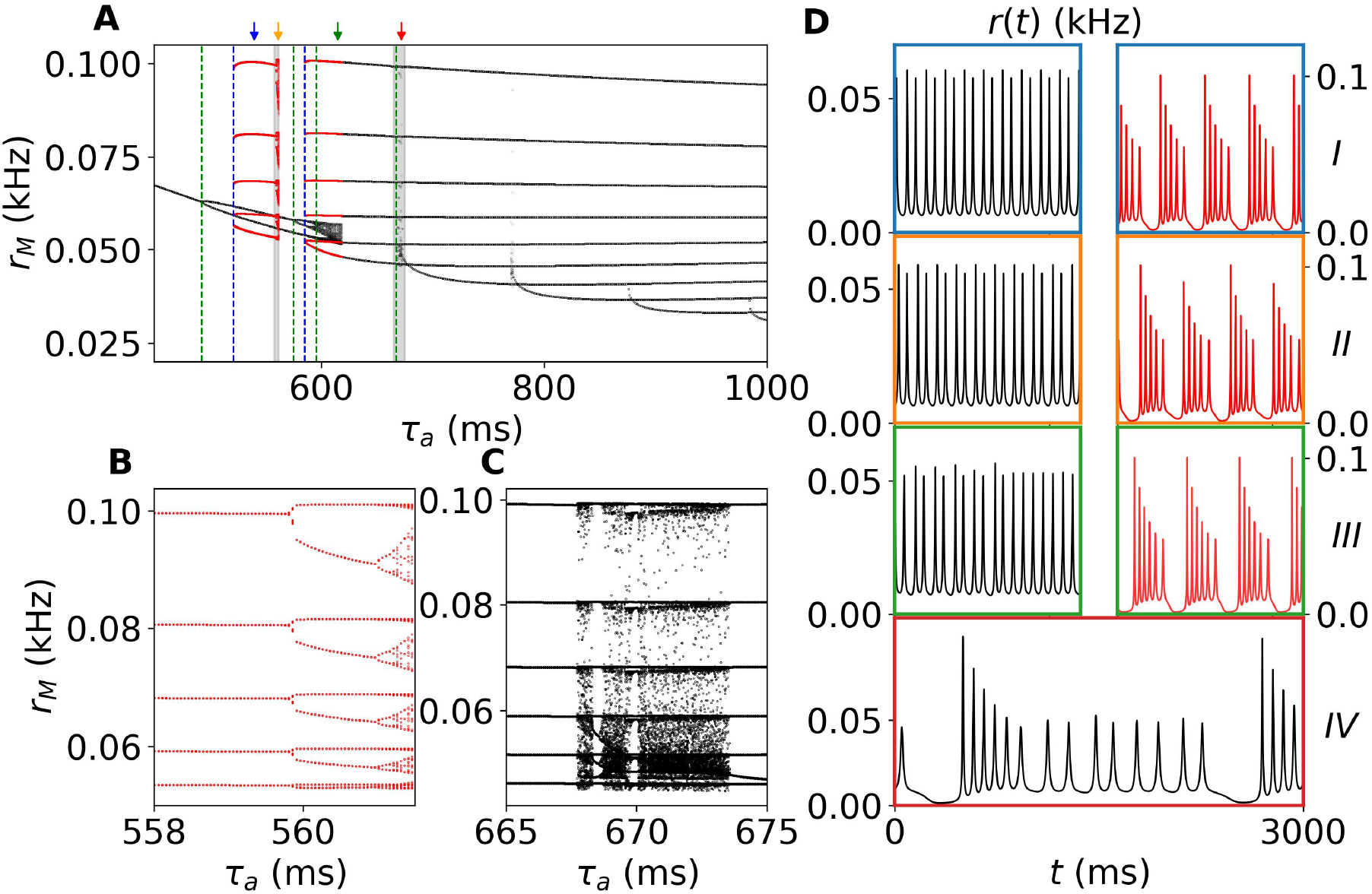
Emergence of population spiking and bursting regimes by varying the adaptation scale *τ*_*a*_. (A) Local maxima of the firing rate *r*_*M*_ obtained via adiabatic continuation in *τ*_*a*_, by both increasing and decreasing its value. At each step, an initial transient of 50 s is discarded, and local maxima are computed over a subsequent interval of 20 s. Numerical integration is performed using an Adams/BDF method with automatic stiffness detection, allowing for accurate resolution of the fast transients arising during bursting dynamics. Both relative and absolute tolerances are set to 10^−12^. Red dots indicate the coexistence of attractors. Green (blue) vertical dashed lines denote period-doubling (saddle-node) bifurcations, as discussed in the main text. Colored arrows highlight selected values of *τ*_*a*_ used in panel (D) for representative time traces. (B) and (C) Zoomed-in views of the local maxima map, corresponding to the gray shaded regions in panel (A). (D) Time traces for selected values of *τ*_*a*_: *I*) *τ*_*a*_ = 540 ms, *II*) *τ*_*a*_ = 561.5 ms, *III*) *τ*_*a*_ = 615 ms, and *IV*) *τ*_*a*_ = 672 ms. In sub-panels *I–III*, the two traces correspond to coexisting attractors, using the same color code as in panel (A). All time traces are shown over a 3 s interval. Other parameters: Δ = 0.1, *J*_*EE*_ = 7, 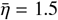, *γ* = 8, *τ*_*m*_ = 20 ms.

For small adaptation time scales (*τ*_*a*_ ≤ 492.82 ms), the system displays periodic PSs characterized by a single peak per cycle. At *τ*_*a*_ ≃ 492.82 ms, a period-doubling (PD) bifurcation gives rise to a two-peak solution with different heights, representing a first spike-adding transition in the PS regime [33]. A second PD bifurcation at *τ*_*a*_ ≃ 575.14 ms generates a three-peak solution and is followed by a cascade of further PD bifurcations. In parallel, a second branch of solutions emerges through a saddle-node bifurcation at *τ*_*a*_ = 521.25 ms. This branch corresponds to ROs whose Up states contain five PBs. The coexistence between the two-peak PS solution and the five-peak RO is illustrated in panel D-I for *τ*_*a*_ = 540 ms. The five-peak RO branch remains stable up to *τ*_*a*_ = 559.627 ms, where it undergoes a PD bifurcation.

The dynamics arising from this bifurcation is particularly unusual. As shown in panel B, the resulting chaotic dynamics after the PD cascades does not modify the number of peaks within each burst. Instead, the amplitudes of successive relaxation cycles become irregular. A representative chaotic trajectory is shown in panel D-II for *τ*_*a*_ = 561.5 ms. Throughout this regime, each burst still contains five peaks, while only their amplitudes are irregular and vary from cycle to cycle.

Returning to the PS branch, the cascade of PD bifurcations eventually leads to a chaotic attractor in the interval *τ*_*a*_ ∈ [590, 619] ms. This regime is characterized by population spikes with irregular amplitudes, as illustrated by the black trace in panel D-III. By analogy with the spiking chaos observed in the Hindmarsh–Rose neuron [33], we refer to this regime as *population spiking chaos*. Interestingly, this chaotic attractor coexists with a second RO branch that emerges through a saddle-node bifurcation at *τ*_*a*_ = 585.13 ms. This branch is characterized by six peaks in the Up state and is represented by the red trace in panel D-III for *τ*_*a*_ = 615 ms. At *τ*_*a*_ = 619 ms the chaotic PS attractor disappears abruptly, suggesting a boundary crisis [12], while the six-peak RO branch persists.

The six-peak RO remains stable up to *τ*_*a*_ = 667.45 ms, where it undergoes another PD bifurcation. The subsequent dynamics is summarized in panel C. In contrast to the chaotic regime associated with the five-peak RO branch, the resulting attractor now exhibits relaxation oscillations containing different numbers of peaks from cycle to cycle. A representative trajectory is shown in panel D-IV. By analogy with bursting chaos observed in single-neuron models [33], we refer to this regime as *population bursting chaos*.

For even larger adaptation times, the system undergoes a sequence of spike-adding transitions. Each transition increases by one the number of peaks contained in the Up state of the stable RO. Consequently, the number of peaks grows progressively, reaching 21 at *τ*_*a*_ = 2400 ms. Successive periodic branches are separated by narrow chaotic windows, which may arise through period-doubling cascades or crisis events. Overall, increasing *τ*_*a*_ leads to a recurrent scenario in which an *n*-peak RO loses stability, passes through regular and chaotic intermediate regimes, and eventually gives way to a stable (*n* + 1)-peak RO.

This phenomenology suggests a chaos-induced spike-adding mechanism: the extra spike is not incorporated through a smooth deformation of the original periodic orbit but rather through an intermediate chaotic transition region. In this sense, the observed dynamics resembles the discontinuous spike-adding scenario described in [34], where additional spikes may arise through chaotic windows, small islands of bursting periodic orbits, and cascades of period-doubling bifurcations, rather than through a purely canard-mediated continuation of periodic solutions.

Since large adaptation time scales give rise to relaxation oscillations (ROs) with different numbers of peaks in the Up state and different oscillation periods, we focus on the representative case *τ*_*a*_ = 2000 ms. The resulting dynamics is characterized in terms of two observables: the period of the RO and the number of peaks occurring during each oscillatory cycle. The corresponding phase diagrams in the 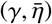 plane are shown in Fig. 5(A-B). A clear spike-adding structure emerges from these diagrams. For fixed 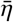, decreasing the adaptation strength *γ* leads to a sequence of transitions in which the number of peaks contained in the Up state increases by one at a time, while the overall period of the RO varies only moderately. This behavior represents a second spike-adding mechanism driven by the adaptation parameters. In contrast to the scenario discussed previously, where spike adding was induced by changes in the adaptation timescale *τ*_*a*_, here the transitions are controlled by the adaptation strength *γ*. As we show below, however, the underlying dynamical mechanisms display important differences.

**Figure 5:**
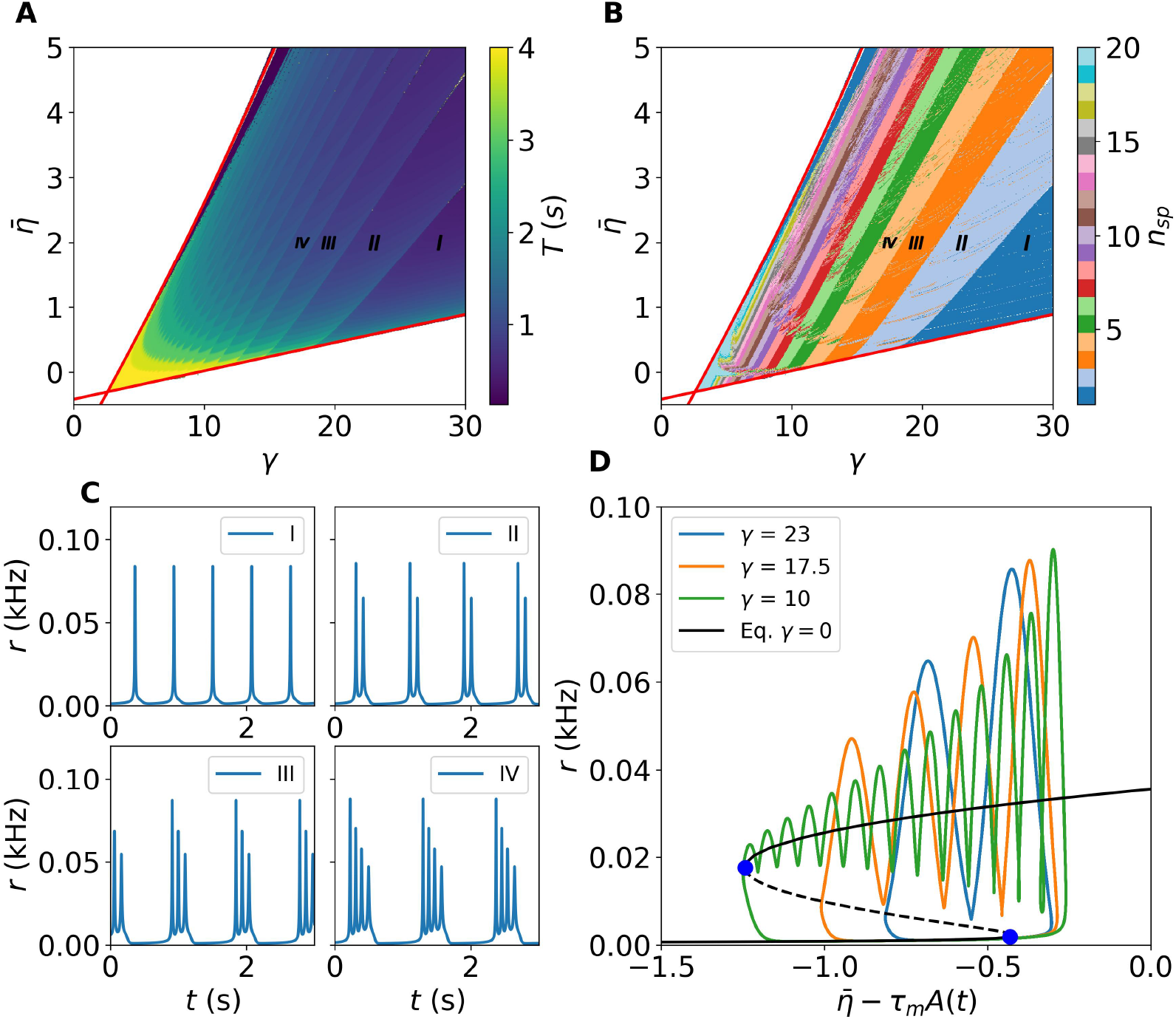
Slow Relaxation Oscillations with Different Number of Peaks. (A) Period of the overall oscillations in the 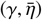 plane. (B) Number of PSs during the Up Phase of a RO in the 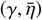 space. Roman numerals identify the regions where are performed the simulations shown in the sub-panels (C). The roman numerals also identify the increasingly added spikes within the identified regions (I-IV). (D) Slow-fast presentation of relaxation oscillations with different numbers of peaks. Black solid and dashed branches stand for stable and unstable fixed points of the FS, respectively. The equilibrium continuation (black curve) as a function of 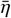 shows two saddle node bifurcations (blue dots). As x-axis we use the input current change due to the fatigue level: 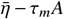. On top we plot the solution of *r*(*t*) as a function of the effective input received 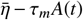 for different values of *γ* identified with the color code. Parameters values are Δ = 0.1, *J*_*EE*_ = 7, *τ*_*m*_ = 20 ms and *τ*_*a*_ = 2000 ms.

The spike-adding sequence can be appreciated in the representative time traces shown in Fig. 5(C), corresponding to regions where the Up state contains 1, 2, 3, and 4 peaks, respectively. As discussed previously, the ROs arise from the interplay between the fast spiking subsystem and the slow adaptation variable. Their evolution can be conveniently visualized in the phase plane 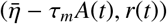, where the RO trajectories of the full system are shown together with the fixed points of the fast subsystem (Fig. 5(D)).

The main effect of varying *γ* is to modify the extent to which the trajectory explores the upper stable branch of the fast subsystem. For large values of *γ*, the strong adaptation-induced self-inhibition rapidly suppresses the population activity, causing the trajectory to leave the high-activity state after only a few damped oscillations and before reaching the saddle-node point. As *γ* is decreased, the effective self-inhibition weakens and the trajectory remains longer on the upper branch, approaching progressively closer to the saddle-node bifurcation. This extended excursion allows additional peaks to develop during the relaxation towards the HAS, thereby generating the observed spike-adding sequence. For sufficiently small values of *γ*, the trajectory follows almost the entire upper stable branch before returning to the LAS (green curve in Fig. 5(D)).

A more detailed characterization of the bifurcations that occur for a fixed value of 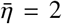, by varying the adaptation strength *γ*, is presented in Fig. 6. In particular, panel (A) shows the maxima of the firing rate *r*_*M*_ obtained by performing quasi-adiabatic simulations: the *γ* values are increased (red) and decreased (black), following the attractor. ^3^ Also in the present bifurcation diagram, regions of coexistence of different solutions are observable. The first coexistence region can be found for small adaptation strengths (see the shaded area on the left around *γ* ∈ (8.9, 9.0)). An enlargement of such region is shown in Fig. 6 (B), where is reported the coexistence of PS solutions (red curves) and a RO with 20 peaks in the Up state. The single PS bifurcates at *γ* ≃ 8.931 giving rise to a solution characterized by 2 PSs of different amplitude, then, at *γ* ≃ 8.962, a third PS of different amplitude adds to the solution. The latter solution leads, via a period-doubling cascade, to a regime of population spiking chaos, characterized by PSs with irregular amplitudes. A characterization of the coexisting solutions in terms of the first two Lyapunov exponents is reported in Fig. 6 (C): one of the two LEs is always identically zero, since we are considering a system of ODEs with continuous time, the other LE becomes zero for the PS solutions (red curve) at each period doubling bifurcation and then definitely positive in the chaotic region. For the RO instead, the second LE (black curve) is always definitely negative in the whole considered range. The abrupt disappearance of the chaotic attractor at *γ* ≃ 8.979 is analogous to the one reported previously and a probable indication of a crisis that leads the system to jump to a RO solution with 19 peaks.

**Figure 6:**
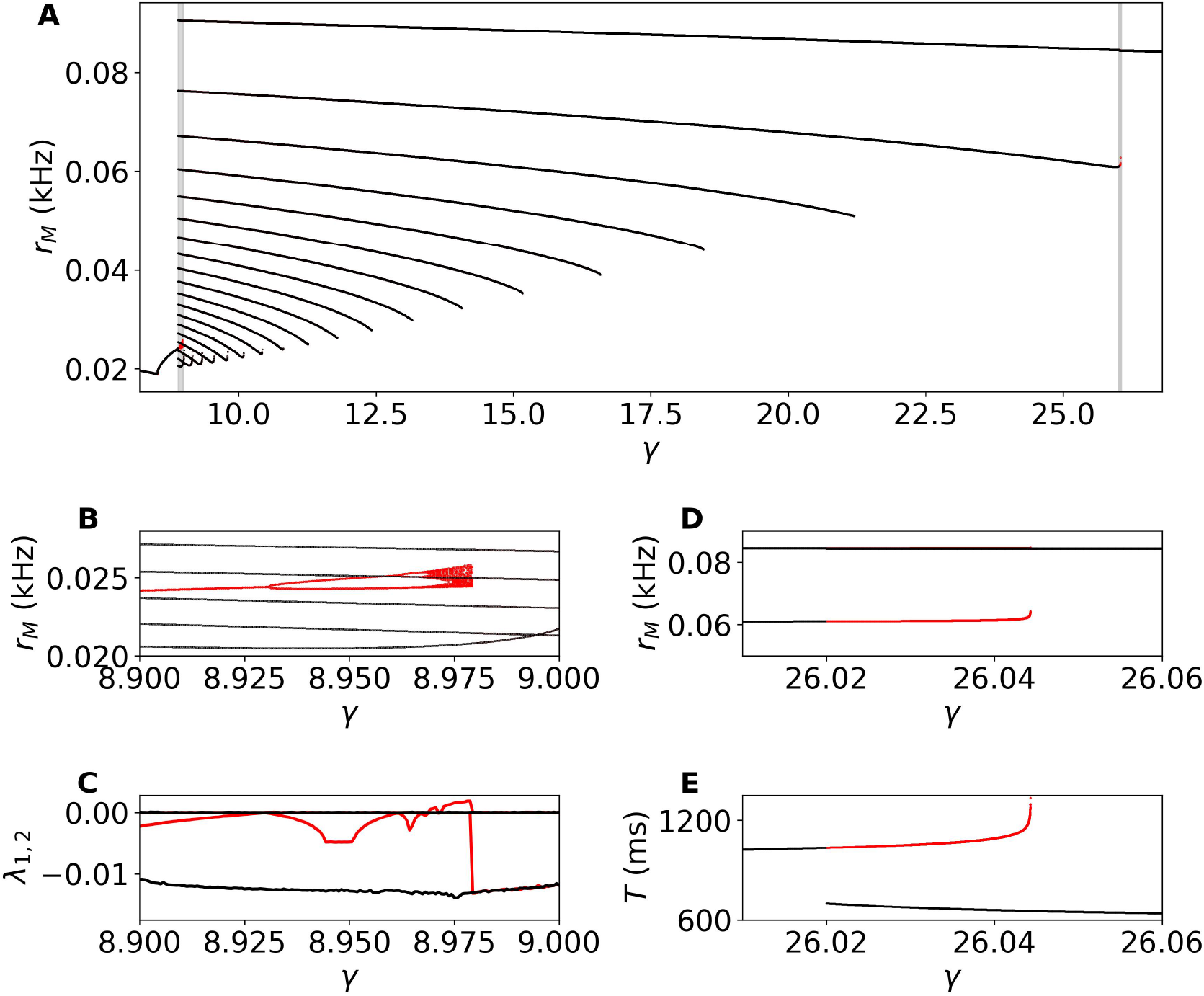
Emergence of population spiking and bursting regimes by varying the adaptation strength *γ*. (A) Local maxima of the firing rate *r*_*M*_ obtained via adiabatic continuation in *γ*. Black (red) dots are the results of the simulations performed by decreasing (increasing) adiabatically the parameter *γ*. The adiabatic results for increasing (decreasing) *γ* are identical except in small parameter regions, that are highlighted in panels (B) and (D). (C) First two maximal Lyapunov exponents *λ*_1_ and *λ*_2_ calculated in the parameter region reported panel in (B). The results obtained by adiabatically decreasing the *γ* parameter reveal that all the solutions are periodic, since *λ*_1_ = 0 and *λ*_2_ *<* 0, instead by adiabatically increasing *γ* one observes a cascade of period doublings (whenever *λ*_1_ = *λ*_2_ = 0) followed by chaos where *λ*_1_ *>* 0. (E) Periods of the RO calculated for the parameter region reported in panel (D). Parameters values are 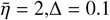, *J*_*EE*_ = 7, *τ*_*m*_ = 20 ms and *τ*_*a*_ = 2000 ms.

In the range *γ* ∈ [8.979; 27.000] we have no coexistence of solutions, and we have been able to identify only Ros with a variable number of peaks in the Up state: the number of peaks increases for decreasing *γ*. Therefore we start now from *γ* = 27 where we have one peak-solution and we adiabatically decrease *γ*. The system now displays a series of spike-adding transitions leading to ROs with a high number of peaks in the Up state: we pass from one peak at *γ* = 27 up to 20 peaks at *γ* = 8.979.

In contrast to the transitions shown in Fig. 4, here we can follow the spike-adding mechanism already starting from the 1-peak solution. This allows for a more detailed characterization of the underlying bifurcation. Indeed, the spike-adding transition for ROs with 1 to 2 peaks—see the zoomed gray region around *γ* ≈ 26—reveals that the first transition occurs via a period-doubling bifurcation at *γ*_1,2_ ≃ 26.04432. At this point, a new stable 2-peaks solution emerges with a period exactly twice that of the 1-peak solution (*T* ≈ 665 for one peak, *T* ≈ 1330 for two peaks), as shown in Fig. 6 (E). Interestingly, this period-doubled solution rapidly decreases its period to a value closer to *T* ≈ 1100 ms, as *γ* is slightly reduced. This quite rapid drop of the period over a very narrow parameter interval renders difficult, for standard continuation methods, to accurately detect the bifurcation. Another notable feature, which departs from the typical period-doubling scenario, is that the period-1 solution does not immediately lose stability. Instead it persists, giving rise to a narrow coexistence region of the 1-peak and the 2-peaks RO solutions in the interval *γ* ∈ [26.02, 26.04].

We have verified that a similar coexistence also occurs for transitions leading from *n*-to (*n* + 1)-peaks ROs. However, the coexistence interval becomes progressively smaller as *n* increases, making it increasingly hard to detect it numerically. Based on this, we hypothesize that the same mechanism operates at each *n* → *n* + 1 spike-adding transition: a period-doubled branch corresponding to the (*n* + 1)-peaks solution emerges and coexists with the *n*-peaks one, with the period doubling bifurcation taking place within an increasingly narrow neighborhood of *γ*_*n,n*+1_. This can be interpreted as a sign of a sub-critical period doubling bifurcation associated with a spike-adding transition, thus resembling the spike-adding transitions described in pancreatic *β*-cells [35]. However, at variance with the results reported in [35], here we do not observe chaotic dynamics in-between the different bursting states, but only in correspondence of the transition leading from population spiking to population bursting shown in panels B and C.

As a last aspect of this analysis and to better visualize the transition from *n*-to (*n* + 1)-spike solutions, we plot the dynamics of 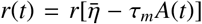 as a function of the control variable 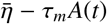 on top of the fixed point solutions (black lines) for the FS. In particular, in Fig. 7 (A), are shown the ROs with 1-peak (blue dots) and 2-peaks (orange line) in the proximity of the bifurcation point *γ*_1,2_. The 1-peak solution shows a closed loop orbit around the lower saddle-node point, delimiting the existence of the LAS in the fast sub-system. The 2-peaks solution emerges at *γ*_1,2_ whenever the orbit does not immediately return to the lower stable branch of the FS, as in the 1-peak solution, but it follows for a while the unstable branch of the FS connecting the two saddle-nodes and then it is attracted back towards the upper stable branch (HAS) of the FS before returning to the LAS. By decreasing further *γ*, the 2-peaks RO detaches progressively from the unstable branch of the FS revealing two damped oscillations around the upper branch before returning to the LAS (see Fig. 7 (B)).

**Figure 7:**
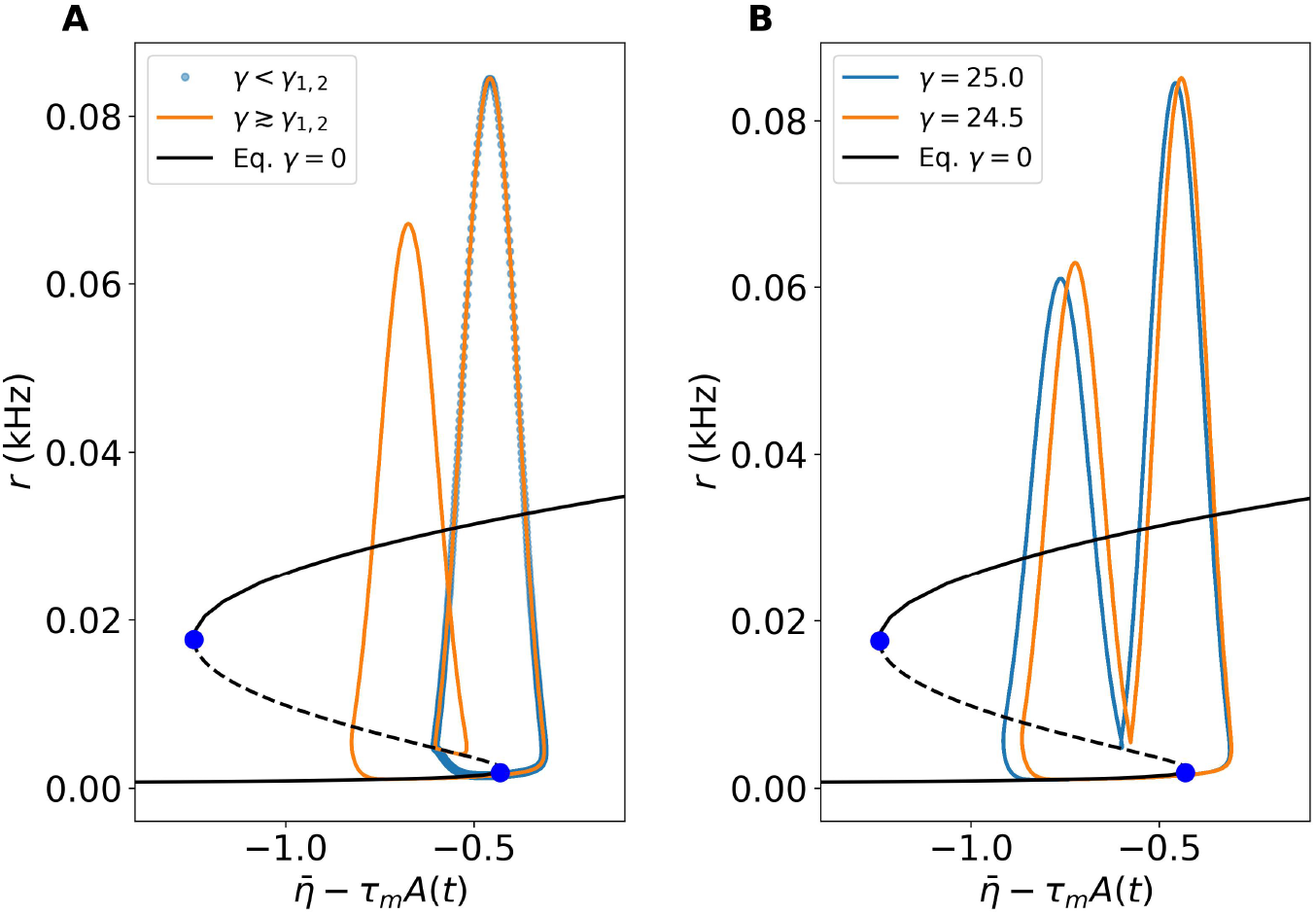
Transition mechanism for the spike-adding transition from *n*-to (*n* + 1)-peaks. The values of the parameter *γ* in (A) are for 1-peak solution (blue dots) *γ* = 26.04431342207, and for the 2-peaks RO (orange line) *γ* ≃ *γ*_1,2_ = 26.04431342206. The black solid (dashed) lines refer to stable (unstable) fixed point solutions of the FS and blue dots to the corresponding saddle-node bifurcations. Other parameters are 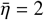, Δ = 0.1, *J* = 7, *τ*_*m*_ = 20 ms and *τ*_*a*_ = 2 sec.

### 3.2. Relaxation oscillations in excitatory-inhibitory networks with SFA

We will now focus on excitatory-inbitory neural mass models with SFA acting on the excitatory population as described in Eqs (9) and (13). For simplicity we will limit to considering the case of symmetric cross-coupling, with *J*_*EI*_ = *J*_*IE*_ = *J*_*C*_, and in the absence of self-inhibitory coupling, i.e. we usually set *J*_*II*_ = 0, unless otherwise stated. The analysis reported in this sub-section is mostly devoted to the characterization of ROs, and in particular of the Up and Down states, in order to compare our findings with experimental results concerning brain waves emerging during Anesthesia or Slow Wave Sleep [1]. For this reason, we will not perform detailed bifurcation analysis as the ones reported in the previous section, but we will focus on the role played by the parameters controlling the excitability of the neurons, the coupling strength between the two populations and the SFA on the emerging characteristics of the ROs.

The observed excitatory–inhibitory oscillatory dynamics is analogous to the one associated to the *pyramidal– interneuronal network gamma* (PING) mechanism [36]. A brief burst of activity in the excitatory population recruits the inhibitory neurons, which respond after a short delay *τ*_*d*_. The resulting inhibitory feedback suppresses the excitatory activity, driving the population into a quiescent state. As inhibition subsequently decays, the excitatory population can become active again, initiating a new oscillatory cycle. In the PS, PB, and chaotic regimes, the inhibitory population is consistently activated after a short delay *τ*_*d*_ following the excitatory activity. The estimated delay lies in the range 3–12 ms, in agreement with the PING mechanism discussed above (Fig. 8).

**Figure 8:**
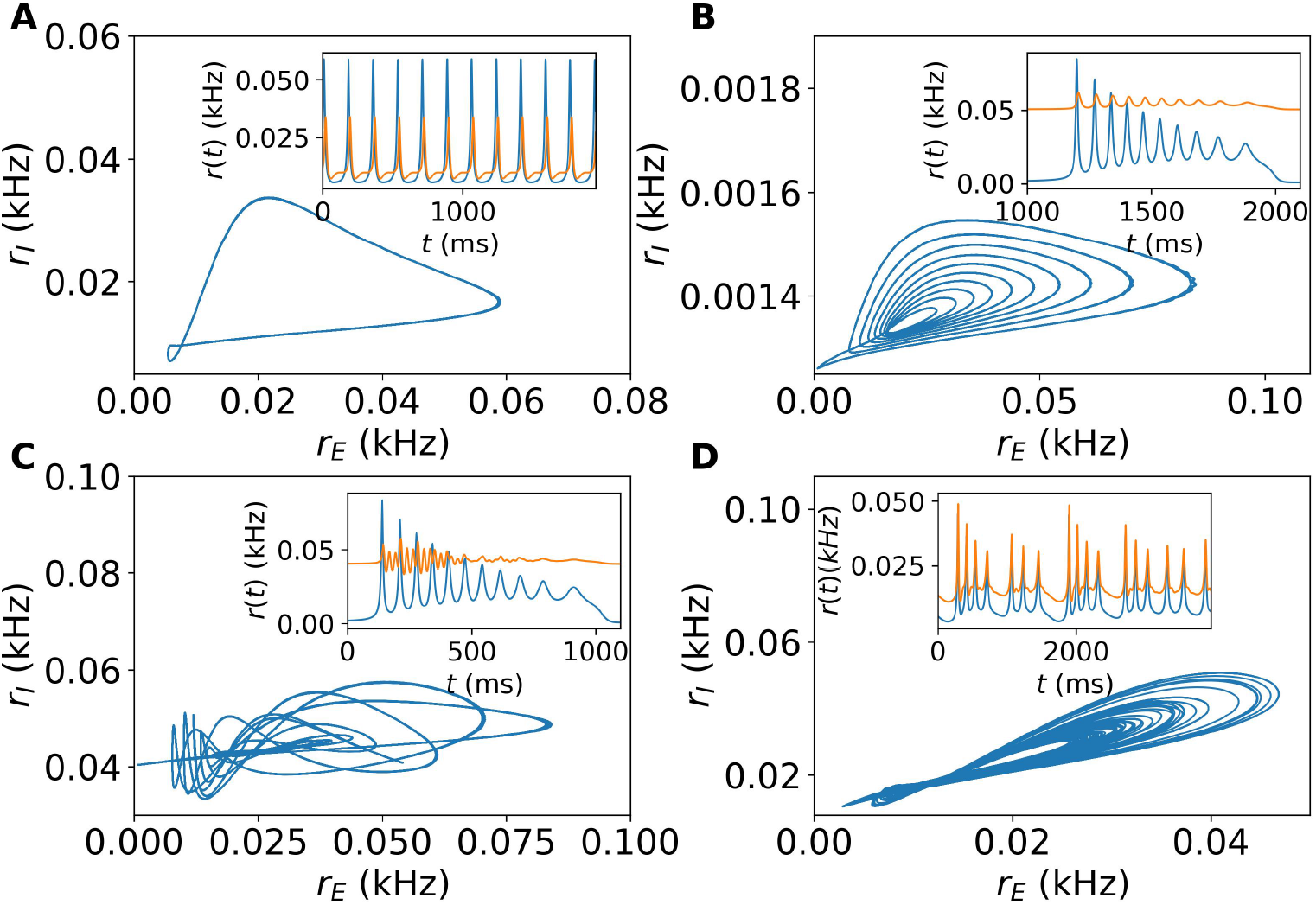
Different dynamical behaviors in excitatory-inhibitory networks. In all panels we show the projection of the attractor in the plane (*r*_*E*_, *r*_*I*_) for different combinations of parameters showcasing the rich dynamics encountered in excitatory-inhibitory neural masses together with the time series of the firing rates for the excitatory (blue) and inhibitory (orange) populations. Both for (A) Population Spiking (*γ* = 6, *J*_*c*_ = 1, 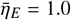 and 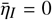) and (B) Population Bursting, we observe a Slow Relaxation Oscillation in the excitatory population (*γ* = 9, *J*_*c*_ = 1, 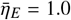 and 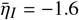). In the inset of this panel we have scaled the inhibitory firing rate by a factor 40 to compare with the excitatory population (C) Slow Relaxation Oscillation in the excitatory population joined to seemingly irregular bursting, but deterministic, in the inhibitory population. (*γ* = 9, *J*_*c*_ = 1, 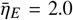 and 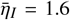) (D) Population bursting chaos (*γ* = 9, *J*_*c*_ = 3, 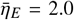 and 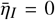). Other parameters 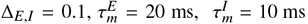, *J*_*EE*_ = 7, *J*_*II*_ = 0, and *τ*_*a*_ = 2000 ms.

While PBs are always associated with slow relaxation oscillations in the excitatory population, the dynamics of the inhibitory population depends strongly on the value of 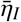. For 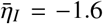, the inhibitory neurons remain mostly below threshold and display oscillations that are largely slaved to the excitatory activity, as shown in Fig. 8(B). A qualitatively different scenario emerges for 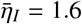 (Fig. 8(C)), where the inhibitory population is intrinsically highly active. In this case, the first excitatory population spike triggers a transient response consisting of damped collective oscillations toward a high-activity equilibrium. However, before this equilibrium is reached, subsequent excitatory spikes repeatedly perturb the inhibitory dynamics, generating a complex yet deterministic temporal pattern. Only after the excitatory burst has terminated does the inhibitory population finally relax toward its stationary HAS.

The coexistence of the intrinsic inhibitory dynamics with the recurrent excitatory drive can give rise to more complex dynamical regimes. In particular, we observe both population bursting chaos and population spiking chaos. An example of the former is reported in Fig. 8(D). These chaotic states typically emerge for sufficiently large coupling strengths (*J*_*c*_ ≳ 3) and excitatory drives 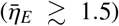, particularly when the inhibitory population operates close to threshold 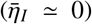. Interestingly, the corresponding attractors remain low-dimensional, with a Kaplan–Yorke dimension only slightly larger than two (*D*_*KY*_ ≈ 2.12).

We now examine the dynamical regimes in the 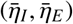 plane for different values of the adaptation strength *γ* and of the excitatory–inhibitory coupling *J*_*c*_ (Fig. 9). The color code indicates the number of peaks in the excitatory firing rate *r*_*E*_(*t*) during each oscillatory cycle: white denotes stationary solutions, blue corresponds to PS dynamics with a single peak, and the remaining colors identify ROs with multiple peaks in the Up state.

**Figure 9:**
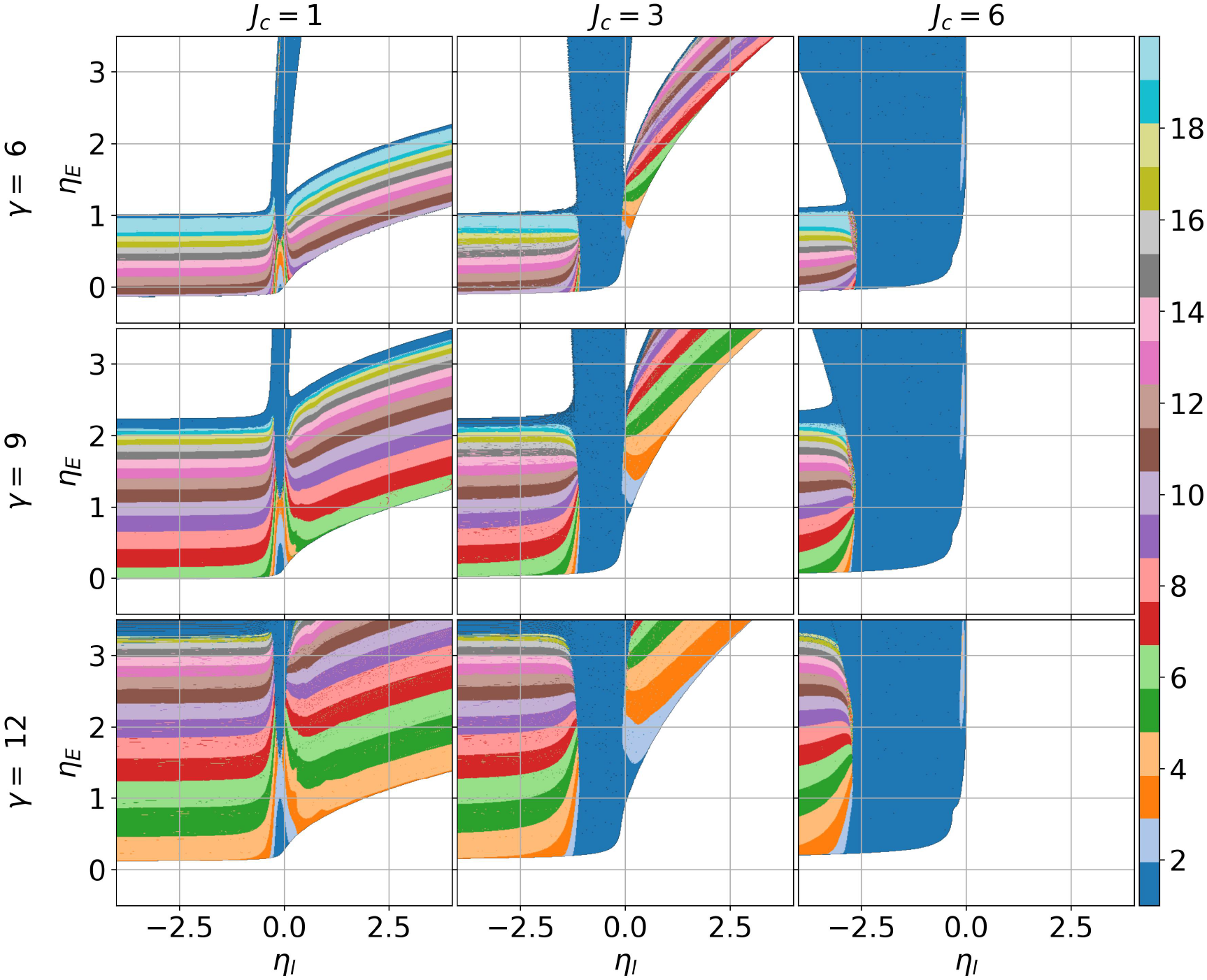
Slow relaxation oscillations for Excitatory-Inibitory Populations. Number of peaks observable during the Up state of the ROs, displayed by the excitatory population firing rate *r*_*E*_ in the 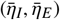 plane for several values of *γ* and *J*_*c*_. Other parameters Δ_*E,I*_ = 0.1, 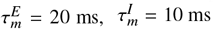, *J*_*EE*_ = 7, *J*_*II*_ = 0, and *τ*_*a*_ = 2000 ms.

A robust spike-adding structure is observed as the mean excitatory drive 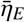 is increased at fixed 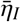. Starting from a LAS, the system enters RO regimes with progressively larger numbers of peaks in the Up state. This is followed by a PS regime and, for sufficiently large 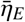, by a HAS. This progression closely resembles the spike-adding scenario found in the purely excitatory network (Fig. 5(B)).

The effect of the inhibitory drive 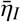 is more subtle. For sufficiently negative 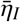, where the inhibitory population is mostly subthreshold, changing 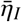 has little effect on the number of peaks in the excitatory ROs. In this region, the colored bands are approximately horizontal, indicating that the spike-adding sequence is mainly controlled by the excitatory drive. Nevertheless, 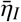 can still affect the temporal structure of the oscillations, in particular the relative durations of the Up and Down states, as discussed below.

In general we observe that around 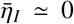 the ROs are usually substituted by a PS dynamics for small cross-coupling (*J*_*c*_ = 1); this region of population spiking enlarges dramatically by increasing *J*_*c*_. Indeed, for *J*_*c*_ = 6 the ROs are present only for definitely negative values of 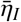, while, as soon as 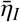 becomes positive, one essentially observes asynchronous dynamics only. This indicates that, for large *J*_*c*_, the inhibitory dynamics strongly influence the behaviour of the excitatory population preventing ROs and bursting.

As previously mentioned, for 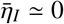, in correspondence of a transition from PS to PB one can also observe chaos. Indeed we have observed, for *J*_*c*_ = 3, population bursting chaos for *γ* = 9 and 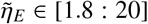 and population spiking chaos for larger values of *γ* = 12 and 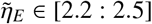.

In the following, we highlight dynamical behaviors of our model that show a qualitative correspondence with recent experimental findings on the characterization of Up and Down states *in vitro*. In particular, Nghiem et al. [1] considered mouse entorhinal cortex slices exhibiting spontaneous slow waves and showed that increasing the carbachol concentration leads to a lengthening of the Up states and a shortening of the Down states. Since carbachol is an agonist of both nicotinic and muscarinic acetylcholine receptors, this pharmacological manipulation can be interpreted as effectively reducing the strength of spike-frequency adaptation [37].

The same effect is observable in our model, as shown in Fig. 10: the increase of *γ* leads to shorter (longer) Up (Down) state durations. This is consistent with the idea that SFA acts as a self-inhibition on the excitatory population, thus limiting the duration of the Up state [20]. As already observed for the purely excitatory population, the increase of *γ* reduces the number of peaks in the Up state and favors a faster return to the Down state.

**Figure 10:**
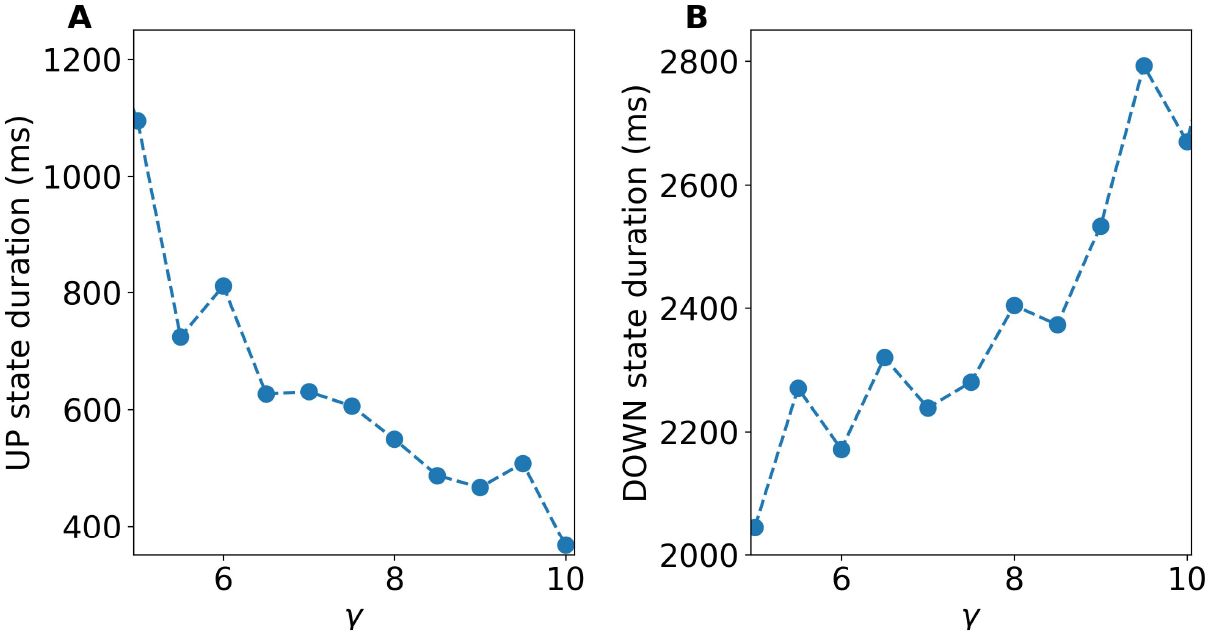
Up and Down state durations are controlled by SFA strength. (A) Variation of Up state duration versus SFA strenght *γ*. (B) Same as in (A) for the Down state. For this figure *J*_*c*_ = 3, *η*_*E*_ = 2.5 and *η*_*I*_ = 1.5. Other parameters as in Fig. 9.

Another important result concerns the correlation between the Down and Up state durations. During anesthesia, it has been reported that the durations of the Down state and the next Up state are positively correlated [21], while, during sleep, the Down and the following Up state durations show a negative correlation. In [1] it has been shown that the increase in carbachol concentration (corresponding to a reduction of the SFA strength) causes a transition from anesthesia-to sleep-like dynamics *in vitro*.

To check whether the same effect is also observable in our model, we fixed all the parameters and considered two different values of the SFA strength, namely *γ* = 9 and 12. The duration of the Up and Down states, obtained by varying 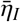 in a limited range, is reported in Fig. 11 (A,B) for the two considered *γ*-values. For *γ* = 12 we observe that the durations of the Down and Up states are clearly correlated with an associated Pearson factor *ρ* ≃ 0.97, while the durations increase when the median inhibitory drive 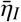 decreases. Furthermore, for *γ* = 9, the Down and Up durations appear as anti-correlated (*ρ* ≃ −0.59); in this case the decrease of 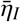 leads to a decrease of the duration of the Up state and to an effective increase of the Down state. Therefore, we can consider the regimes at *γ* = 12 (*γ* = 9) as putative anesthesia-like (sleep-like).

**Figure 11:**
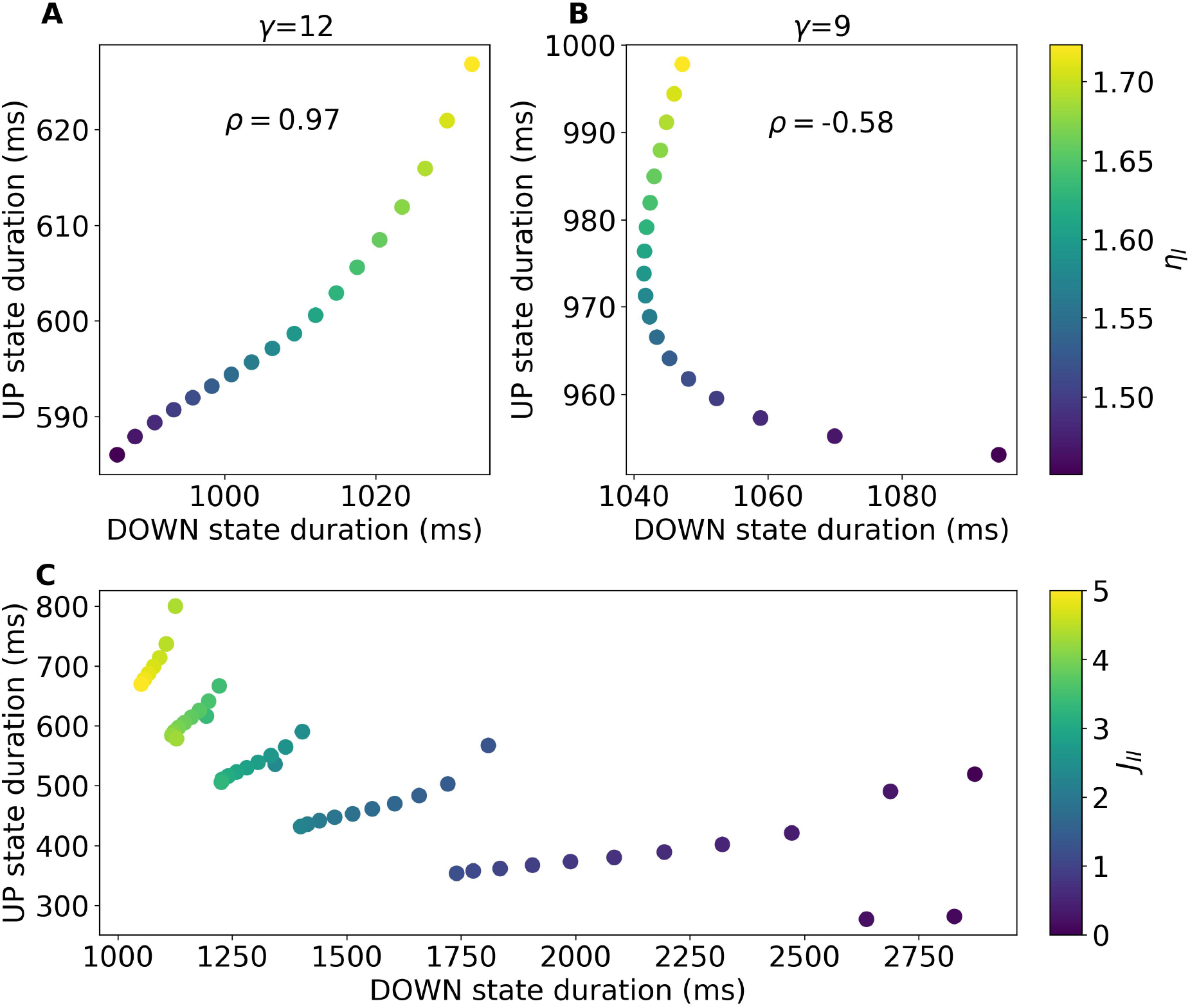
Correlations of Down and Up state durations. EXPERIMENT 1: (A) Up state duration as a function of the Down state duration for several 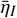 values (shown in the color bar) and fixing 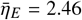, *J*_*c*_ = 1 with *γ* = 12. Text shows the correlation of the data points. (B) Same as in (A) by decreasing SFA strength to *γ* = 9. EXPERIMENT 2: (C) Up state duration as a function of the Down state duration at several values of *J*_*II*_ (shown in the color bar) and fixing 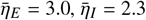, *J*_*c*_ = 3 with *γ* = 9. Other parameters as in Fig. 9

The assumption that the adaptation strength controls the transition from sleep-like (low *γ*) to anesthesia-like (high *γ*) states is somehow confirmed by the data reported in Fig. 10(B), since the increase of the Down state duration passing from slow wave sleep to deep anesthesia has been reported for different cortical areas in cats [38] and for the primary visual cortex in chronically implanted rats [22].

As a final example, we consider the experimental results reported in [25]. In that study, the authors analyzed cortical slices displaying slow waves and investigated how the dynamics was modified by reducing inhibition through the application of gabazine or bicuculline methiodide. One of their main findings was that a progressive reduction of inhibition leads to a shortening of the Up-state duration and a lengthening of the Down state, together with a decrease in the oscillation frequency. This effect was accompanied by a higher firing rate during the Up state and by a longer and deeper hyperpolarization of excitatory neurons. Based on these observations, the authors suggested that an increase in adaptation could play a compensatory role in response to the reduced level of inhibition in the network [25].

To mimic this experiment, we fixed all model parameters and varied only the self-inhibitory coupling *J*_*II*_ in the interval ([0,5]). The results are shown in Fig. 11(C). As *J*_*II*_ decreases, the model reproduces the experimentally observed shortening of the Up state and lengthening of the Down state. Conversely, decreasing *J*_*II*_ leads to a noticeable increase in the period of the RO, from approximately 1.7 s at *J*_*II*_ = 5 to 3.4 s at *J*_*II*_ = 50. As shown in the figure, this overall trend occurs through a sequence of intermediate steps in which the Up- and Down-state durations display opposite dependencies on *J*_*II*_, in qualitative agreement with recent findings reported in [22]. Each step corresponds to a relaxation oscillation with a well-defined number of peaks during the Up state, ranging from 9 peaks at *J*_*II*_ = 5 down to 4 peaks at *J*_*II*_ = 0. Thus, the shortening of the Up state is associated with a reduction in the number of peaks present in each relaxation oscillation. We also observe that decreasing *J*_*II*_ leads to a longer hyperpolarized phase, as measured from the mean membrane potential *v*_*e*_ (data not shown). However, in contrast to the experiment, this effect is accompanied by a decrease, rather than an increase, of the mean excitatory activity during the Up state.

From the analysis of our model the effects of inhibition reduction observed in [25] are reproducible in specific cases only: namely, by reducing the recurrent coupling within the inhibitory population or the activity of inhibitory neurons whenever their dynamics is essentially slaved to the excitatory one.

## 4. Summary and Conclusions

In this paper, we have considered the effect of Spike Frequency Adaptation (SFA) in promoting the emergence of Relaxation Oscillations (ROs) in Next Generation Neural Mass Models (NG-NMMs). First, we considered a single excitatory population subject to a global slow adaptive current that accounts for the fatigue associated with spike emissions. The possible dynamical states are visualized in the phase diagram displayed in Figs. Fig. 2(B), as a function of two main parameters controlling the overall dynamics: the median excitatory drive 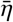 and the adaptation strength *γ*. At low 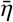, the system displays an asynchronous low-firing state (LAS), whereas at high 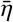 it exhibits an asynchronous HAS. In between these two regimes, collective oscillations emerge within a triangularly shaped region, while for low values of both *γ* and 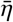, coexistence between the two asynchronous states can be found. This scenario is closely analogous to the one reported in [20, 22] for a network of excitatory-inhibitory Leaky Integrate-and-Fire neurons with SFA acting on the excitatory population.

The adaptation timescale *τ*_*a*_ controls the dynamical evolution of the collective oscillations. In particular, for sufficiently long relaxation times (*τ*_*a*_ *>* 500 ms), one observes the emergence of slow ROs characterized by the periodic alternation of a high-firing state (Up phase) and a low-firing state (Down state). A slow-fast analysis clarifies that the fast dynamics is characterized by the coexistence of a low-firing node and a high-firing focus connected by an unstable saddle [8]. The slow adaptation drives the orbit from the LAS towards a saddle-node, beyond which the trajectory is attracted by the HAS. The orbit then relaxes towards this state through a series of damped oscillations, corresponding to population spikes, before being eventually re-attracted towards the LAS.

For the considered parameters, we observed ROs with a number of peaks ranging from five at *τ*_*a*_ ≃ 500 ms up to 21 peaks at *τ*_*a*_ ≃ 2400 ms. The number of peaks in the Up phase increases with *τ*_*a*_ through a discontinuous spike-adding mechanism, in which additional peaks arise through a sequence of regular and chaotic windows, with intermediate cascades of period-doubling bifurcations. This scenario is similar to the chaos-induced spike-adding mechanism reported in [34] for the Hindmarsh–Rose model.

In this scenario, we observe peculiar situations, such as a period-doubling cascade leading to population spiking chaos, characterized by PSs with different amplitudes. This cascade coexists with stationary ROs and even with chaotic ROs, in which the number of peaks per RO remains fixed, while their amplitudes are irregularly distributed across successive cycles. Furthermore, in correspondence with the spike-adding bifurcations, we also observed population bursting chaos, characterized by sequences of ROs, or population bursts, with different numbers of peaks. The terms population spiking chaos and population bursting chaos are used here by analogy with the spiking and bursting chaos observed in the single-neuron Hindmarsh–Rose model [33]. Moreover, population spiking and population bursting chaotic regimes have also been reported in Kuramoto models with and without inertia in presence of global adaptation [39].

The analysis of the adaptive excitatory neural mass as a function of the adaptation strength *γ*, for a fixed slow adaptation timescale *τ*_*a*_ = 2000 ms, reveals that increasing *γ* at fixed median excitatory drive leads to a drastic decrease in the number of peaks during the Up phase, eventually giving rise to population spiking and even to asynchronous dynamics. Since the effect of adaptation is essentially inhibitory, increasing the adaptation strength amounts to increasing the effective inhibition. This prevents the emergence of damped global oscillations during the Up state and drives the orbit back towards the Down state more rapidly.

The emergence of ROs with an increasing number of peaks, controlled by the parameter *γ*, occurs via a different spike-adding mechanism. We observe that solutions with (*n* + 1) peaks emerge through a period-doubling transition, while coexisting, in a very narrow parameter region, with the RO displaying *n* peaks. Thus, we conjecture that these spike-adding bifurcations resemble those observed in a three-variable model reproducing the characteristic bursting and spiking behavior of pancreatic *β*-cells [35]. Indeed, for that model, the authors showed that spike-adding transitions consist of saddle-node bifurcations in which the (*n* + 1)-spike bursting behavior is born, slightly overlapping with a subcritical period-doubling bifurcation in which the *n*-spike bursting behavior loses stability. However, in [35], chaos tends to arise in the transitions between continuous spiking and bursting, as well as between different bursting states. In contrast, in our case, we identified a chaotic attractor coexisting with ROs only in the proximity of the spiking-bursting transition at low *γ* values, but not in correspondence with the spike-adding bifurcations.

Then, we extended the model by considering excitatory-inhibitory populations with SFA acting on the excitatory population. This model includes the essential ingredients usually employed to mimic the emergence of brain oscillations during anesthesia or Slow Wave Sleep [23, 1, 40, 26]. Also in this case, by considering the firing rate of the excitatory population, we observe the same dynamical states found in the purely excitatory case: namely, asynchronous dynamics, population spiking, relaxation oscillations, and the types of chaos previously discussed. The inhibitory response follows the excitatory stimulation as in a typical PING scenario, with a delay in the response of ≃ 3 −12 ms [36]. Furthermore, the dynamics of the inhibitory population appears to be essentially slaved to that of the excitatory one when inhibitory neurons are mostly subthreshold, i.e., for 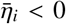. On the other hand, for high levels of inhibitory excitability, 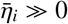, the dynamics of the inhibitory population becomes apparently quite complex. Indeed, after each excitatory PS, the inhibitory population responds by displaying damped oscillations towards a high level of activity; if additional excitatory PSs arrive before the final relaxation, the evolution of the inhibitory firing rate becomes quite intricate, although still not chaotic. Population spiking and population bursting chaos are instead observed for 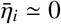, in correspondence with the transition from population spiking to population bursting, or ROs.

The last part of the analysis was devoted to understanding whether this minimalistic model is able to capture, at least qualitatively, some of the experimental results found *in vitro* concerning the characterization of spontaneous slow waves. We first considered the effect of carbachol, an agonist of acetylcholine receptors, which can be interpreted as effectively reducing the strength of spike-frequency adaptation [37]. In [1], it was shown that increasing the carbachol concentration leads to a lengthening of the Up states and a shortening of the Down states, and can also drive a transition from anesthesia-like to sleep-like dynamics. Consistently with this interpretation, in our model a decrease in the adaptation strength *γ* leads to longer Up state durations and shorter Down state durations. Furthermore, by fixing all the parameters in our model and increasing only the adaptation strength we pass from a situation where Down state and Up state durations are anti-correlated (as during sleep) [1] to another one where the duration of the Down and Up states are positively correlated (as during anesthesia) [21, 22].

Finally, we considered the effect of inhibition reduction, induced by the perfusion of gabazine or bicuculline methiodide in slices of visual cortex [25]. We were able to reproduce the negative correlation between Up and Down state durations observed in cortical slices by reducing the strength of the recurrent inhibitory coupling.

These results are encouraging and suggest that our model, despite its extreme simplicity, can be a useful candidate for better understanding the mechanisms controlling neural dynamics during anesthesia and sleep, as well as for simulating brain-level activity with realistic connectomes. One of the main limitations of our model is that it is fully deterministic. Therefore, it cannot reproduce the variability in Up- and Down-state durations usually observed *in vitro* and *in vivo* [1, 22]. A mean-field formulation encompassing both SFA and noisy fluctuations could be envisaged by considering either a generalization of NG-NMMs, as in [41], or a Fokker–Planck formulation, as done in [42]. Furthermore, in the same framework also finite size fluctuations could be eventually self-consistently introduced by applying the approach developed in [43].

Another limitation of the model is that adaptation is considered as a global effect. This issue has been recently addressed in [44], where the authors developed an exact low-dimensional neural mass model for a heterogeneous QIF network with local adaptive currents. However, in order to obtain this remarkable result, they introduced an adaptation current that depends quadratically on the firing activity of each QIF neuron, rather than linearly as in the present case. In the future, it will be interesting to analyze how the emergence of relaxation oscillations is affected by the different forms of SFA.

## Acknowledgments

The authors acknowledge useful discussions with Roberto Barrio, Pau Clusella, Maurizio Mattia, Nina La Miciotta, Francesco Marino, Michele Valla, Gianni V. Vinci.

## Footnotes

2 Quasi-adiabatic simulations reported in Fig. 4 are performed by increasing/decreasing with variable steps depending on the resolution required but typically in the range Δ*τ*_*a*_ = [0.01 : 0.1] ms. At each step a transient of duration 40 sec is discarded, and the estimation of the local maxima is done on an interval of 10 sec duration.

3 Quasi-adiabatic simulations reported in Fig . 6 are performed by increasing/decreasing *γ* with variable steps Δ*γ* depending on the required resolution, but typically in the range Δ*γ* = [1 × 10^−5^ : 1 × 10^−2^]. At each step a transient of duration 40 sec is discarded, and the estimation of the local maxima is done on an interval of 10 sec duration.

